# A discrete-to-continuous mathematical model for ensemble distributions of a ligand-interacting macromolecular species across milieux-dependent conformational states may offer insights into the genesis and progression of cooperative binding

**DOI:** 10.64898/2026.06.26.734722

**Authors:** Siddhartha Kundu

## Abstract

Small molecule modifiers whence bound, allosterically, will alter the binding of a macromolecule to one- or more-cognate substrates/partners via conformational and non-conformational changes. Although allostery is inferred directly from empirical data, the mathematical basis of these models, constraints deployed and choice of parameter(s) are not clear. Here, we present and characterize a discrete-to-continuous mathematical model for ensemble distributions of a ligand-interacting macromolecular species across milieux-dependent conformational states and examine its’ role in the genesis and progression of cooperative binding. The premise, of our model, is a set of occupancy matrices (sparse, binary, strictly delocalized) which can be partitioned by a probability-based hyperparameter into mutually exclusive proper subsets of occupancy matrices with identical multinomial probabilities. Since each subset is canonical with a constituent occupancy matrix, it is characterized by a unique multinomial probability. The inner product of combinatorial pairs of all mutually exclusive subsets of occupancy matrices, with an expression for the summed transitional probabilities (finite differences between unique multinomial probabilities), is the differentiable matrix of strictly positive real-valued numbers for the system of ensemble distributions. Whilst the harmonic mean is presented as a generic solution for a system of ensemble distributions, the row-wise definite integral for each column is the finite union of open intervals (contiguous, strictly monotone) which in tandem with a set of interval-specific and bounded transitional probabilities constitutes a piecewise smooth curve (path-connected-, closed- and compact-set). Our discrete-to-continuous model is phenomenological and able to recapitulate the basic tenets of cooperative binding whilst offering insights into the genesis and progression of the same.

## 1. Introduction

A ligand-macromolecular complex can either be non-enzymatic or facilitate the conversion of one or more substrates into product(s). The nature of this interaction (qualitative, quantitative) determines the stability of the formed complex and is an important determinant of biochemical function [1-6]. Although these interactions are non-covalent (hydrogen bonds, electrostatic, Van der Waals, hydrophobic), transient covalent linkages are not unknown [7-11]. The latter includes the disulfide (−S−S −)linkage between the thiol (−SH) -group of pairs of cysteine residues of reduced Glutathione (G−SH)-monomers and the binding of peptides to ERp57 in the assembly/disassembly of the major histocompatibility complex I (MHC1) [9-11]. Direct alterations to the active-site of an enzyme notwithstanding, catalysis may also be regulated by long-range conformational changes. In order for these to occur and remain efficacious an inherent and concomitant flexibility (rmsd ≫ 1 Ang) of the active-site residues is likely to be complementary, in contradistinction to enzymes with increased stiffness of the same (rmsd ≈ 1 Ang) [12]. For example, catalysis by the peptidyl-dipeptidases such as the angiotensin-converting enzymes (ACEs) and the signal transducing guanosine triphosphatases (GTPases) are considered “promiscuous” in terms of substrate selection and may be attributed to the existence of stable multistate active-site residues [12]. Although conformation-driven changes, in the genesis of allostery, are well documented, the generation and transmissibility thereof, of low-frequency vibrations in the genesis of non-conformational (entropic, dynamic)-allostery cannot be over emphasized [13-15].

Allostery is defined as the effect of non-orthosteric ligand(s) on the binding/disassembly/activity of a macromolecule [16, 17]. The biochemical- and physiological-relevance of the hemoglobins (*HbAA, HbFF, HbSY;Y* = {*A,S*}) and the set of heterotrope ligands (oxygen, carbon monoxide, cyanide, 2,3-Bisphosphoglygerate (BPG)) are archetypal, of allostery, with ramifications for regular (*O*_2_-transport or -storage, acid-base balance, *CO*_2_-exchange)- and disease (sickle cell disease, hemolysis, toxicology)-biology [7, 18-21]. High-affinity peptide binding by the major histocompatibility complex (MHC)-1 complex is a fundamental pre-processing step, by all nucleated cells, prior to antigen presentation, detection and processing by CD8-cytotoxic T-cells [22-24]. Peptide-editing is the dissociation of low-affinity peptides from MHC1 and is mediated by the peptide loading complex (PLC) and may be driven by low-affinity peptides as part of the retrograde transport pathway from the ER [22-24]. The binding of small molecules (substrate, activator, inhibitor) to an enzyme, at one or more allosteric- or orthosteric-sites, will result in modification of its catalytic (specific)-activity. This can improve/reverse catalytic turnover and/or substrate binding (cytosolic 5’-nucleotidase II, EC 3.1.3.5; Glyceraldehyde-3-phosphate dehydrogenase, EC 1.x.y.z) [25, 26] or both as in the ping pong mechanism (double displacement) of transaminases (EC 2.x.y.z), ordered single displacement (NAD(P)H-dependent oxidoreductases, EC 1.x.y.z) and the random single displacement (kinases, EC 2.x.y.z) [27-30]. Conversely, the identification of novel allosteric sites on macromolecules may facilitate the refinement of potential pharmacophore(s) from libraries of ligands [31, 32]. Empirical data, in tandem with theoretical studies, affirms that allostery-mediated changes are ill-defined (long-range conformational, low-frequency vibrational) and will contribute to the integrity and stability of the ligand-interacting macromolecular complex [6, 13-15, 33-47]. Newer applications for allostery-based regulation include homotrope-multivalency in the N-Acetylgalactosamine (GalNAc)-mediated delivery of siRNA to the liver (ligand-mediated RNAi) and mapping for novel allostery-facilitating amino acids (site-directed mutagenesis) [31, 32, 48, 49].

As alluded to, vide supra, a fundamental outcome of allostery is cooperative binding/unbinding which is described as the effect of a predecessor ligand on the binding of a successor ligand. In other words, whilst “positive”-cooperativity will occur when a predecessor ligand(s) facilitates the binding of successor ligand(s), “negative”-cooperativity is the accelerated disengagement of successive ligands by the binding of a predecessor ligand(s). The operatic molecular mechanisms include intra- and inter-molecular interactions, long range conformational changes, low-frequency vibrations and thermodynamic fluctuations [11-15, 22, 41-43]. The concerted model of Monod, Wyman and Changeux (MWC) posits the existence of distinct activated (relaxed, R; open, O)- and inhibited (taut, T; closed, C) conformations for an allosterically-regulated enzyme/protein [50]. The transient dynamics of the ligand interaction (bound ⇌ unbound) with a single subunit will induce a conformational change which is shared with other subunits. This will result in an alteration in the strength-of-binding, shift in the R or O ⇌ T or C equilibrium and altered dynamics for the cognate substrate(s)/partner(s) [50]. The MWC-model is a reliable representative of cooperative binding in homotropic systems such as the binding of upto four molecules of molecular dioxygen binding to Hemoglobin and the N-succinylamino acid racemases (EC 4.2.1.113) [12, 50, 51]. The sequential model of Koshland, Nemethy and Filmer (KNF) suggests that individual subunits need not possess identical conformations and that ligand binding alone will introduce subtle long range conformational changes so as to facilitate/restrict the binding of similar (homotrope)- or dissimilar (heterotrope)-ligands to other subunits [52]. This model is able to reproduce the kinetic data for a heterotropic system of ligands (oxygen, carbon dioxide, 2,3-BPG) for Hemoglobin [52].

Unlike the MWC- and KNF-models which are strictly empirical, later models are phenomenological and have attempted to incorporate empirical data in tandem with hyperparameters for noise, errors and variance estimation [12, 53-55]. The “Morpheein” model for allostery posits the presence of finite, equally probable, thermodynamically stable and differentially active, core structural conformations (n > 2) in dynamic equilibrium with each other [53-55]. These higher-order distinct conformers are formed by deploying non-covalent interactions and includes catalysis by Porphobilinogen synthase (n ≥ 3,EC 4.2.1.24), Pyruvate kinase (n ≥ 5, EC 2.7.1.40), Ribonuclease A (n ≥ 6, EC 3.1.27.5) Malic enzyme (n ≥ 4, EC 1.1.1.40) along with non-catalytic (Tumor necrosis factor-alpha) proteins [56-62]. These data suggest that cooperative binding/unbinding and the ensuing kinetics are functions of low- and medium-level fluctuations of several macromolecular residues (orthosteric, allosteric). The unified ensemble framework extends these notions by combining a thermodynamically averaged-structure with principles of information theory in an energy landscape model [43-46, 54, 55]. For example, Alanine, a hydrophobic and dietary non-essential amino acid with an approximate MW of 89 g/mol, is often utilized in site-directed mutagenesis studies (combinatorial Alanine scanning) as a rapid screen to identify ‘hot’-spots in the energy landscape of protein(s)-of-interest whilst concomitantly assessing the reactivity for exogenous small molecules, thereby reprising its role as functional sink (inert, structurally non-disruptive) [12, 43, 59-62].

The preponderance of empirical models, for allostery, notwithstanding, the consideration of thermodynamic principles and inclusion of hyperparameters that result thereof, as descriptors, into numerical expressions that describe these have gained considerable traction [13-15, 48-62]. However, the mathematical basis of these models are either understated, presumptive or subsumed entirely. Here, we present a discrete-to-continuous model to comprehend the genesis, progression and biochemical relevance of cooperative binding for ensemble distributions of a ligand-interacting macromolecular species across milieux-dependent conformational states. In section 2 we outline and develop the model with definitions, lemmas, theorems and corollaries, wherever applicable. In “Results” (Section 3), mathematical analyses of the model will be presented and discussed. Section 4 will highlight the biological relevance of our mathematical model by modelling the oxygen dissociation curve (ODC) for Hemoglobin and some of its variants (non-catalytic) and the kinetics of Aspartate transcarbomylase (ATCase)-mediated catalysis of L-Aspartate. Concluding remarks (Section 5) will include a summary of the presented study, limitations and future directions. Mathematically pertinent abbreviations include “D” (definition environment), “L” (lemmas), “T” (theorems) and “C” (corollaries). Section 6, includes detailed proofs for most of the aforementioned results.

## 2. Notation, definitions and preliminary results

In this section we present biochemically relevant paradigms which we utilize to develop our discrete-to-continuous model for ensemble distributions of ligand-interacting macromolecular species that are distributed across milieux-dependent conformational states.

### 2.1 Definitions pertinent to developing a discrete-to-continuous mathematical model for ensemble distributions of ligand-interacting macromolecules

We define a “conformational”-state as the set of prevailing physicochemical- and biochemical-conditions that is likely to exist if and only if a finite number of identical macromolecules adopts the same conformation (***Def. (1)***). We describe the same with a random number and probability of occurrence (*P*(·) = 1) (***Def. (2)***). Paraphrasing these statements,

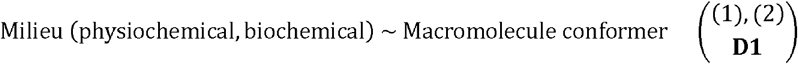

This definition, although simple, allows us to model and thence equate numerous variables that span both, the local microenvironment (physicochemical, biochemical) and the structural descriptors that characterize a macromolecule and justifies the annotation as a definition environment (**D**). **D1** also implies the existence of a countably infinite number of *a*-indexed A-conformational states (x_*a*=1_, …, x_*a*=*A*_); ***Eq***. (**3**)) which we model as a vector of independent and identically distributed random numbers,

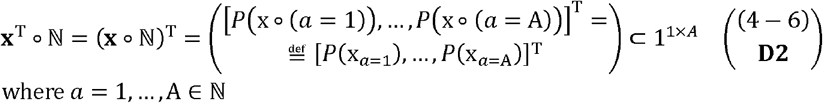

We denote a finite number of *i*-indexed identical and ligand-interacting *M*-macromolecules (*m*_*i*=1_, …, *m*_*i*=*M*_); ***Eq***. (**7**)) as a “species”, which we model as a vector of independent and identically distributed random numbers (***Def. (3)***),

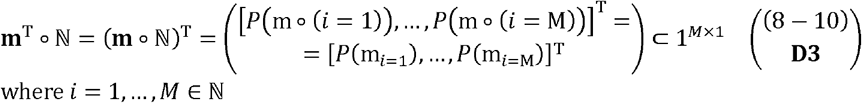

On the basis of **D1**-**D3**, we define the “occupancy” of the *a*^*th*^-conformational state by the *i*^*th*^ -ligand-interacting macromolecule, with the intersection operator and compute the joint probability,

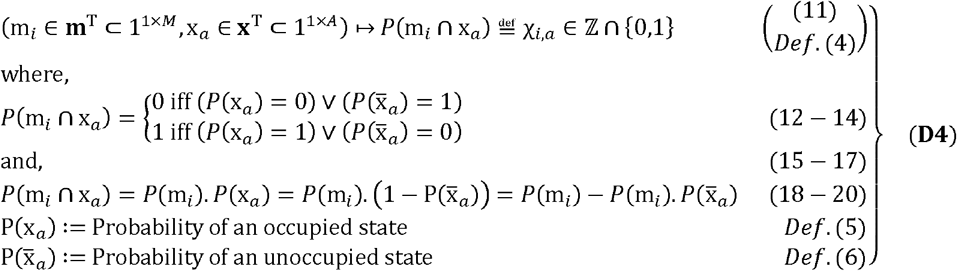

#### Corollary 1

(**C1; without proof**):

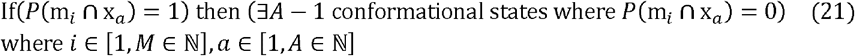

#### Lemma 1

(**L1**; Section 6, **Eqs. (1-14)**):

The finite conformational states across which a ligand-interacting macromolecular species is distributed can be expressed as linearly independent components of a unit or elementary vector,

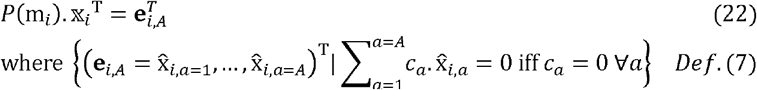

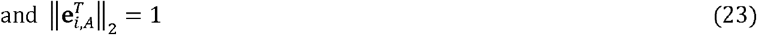

The intracellular- and extracellular-microenvironments that macromolecules are likely to interface with are dynamic, complex and comprise molecular- and non-molecular-components. This means that the concurrent clustering of all members of a macromolecular species at any single conformational state is an improbable biochemical event. We also subsume, as finite, the cardinalities for the ensemble distribution (ligand-interacting macromolecular species, conformational states),

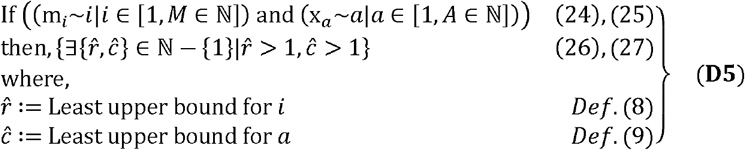

With these definitions and results we define and utilize, hereafter, the finite- and ordered-pair 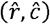 to describe an ensemble distribution of ligand-interacting macromolecular species across conformational states,

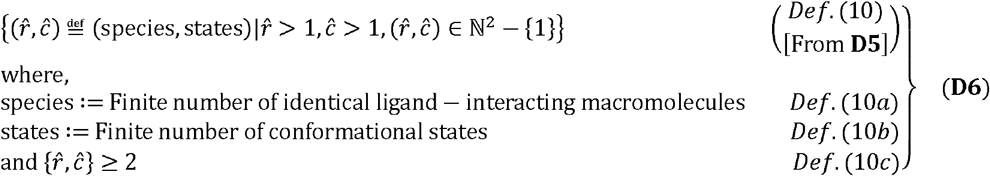

### 2.2 Occupancy matrix, for an ensemble distribution, as the basis of our discrete-to-continuous mathematical model

We now describe the construction, characterization and parameterization of an occupancy matrix, for an ensemble distribution.

#### Corollary 2

(**C2;** Section 6, **Eqs. (15-20)**):

The occupancy matrix for an ensemble distribution is,

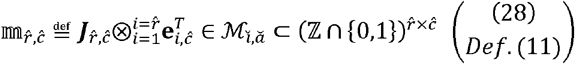

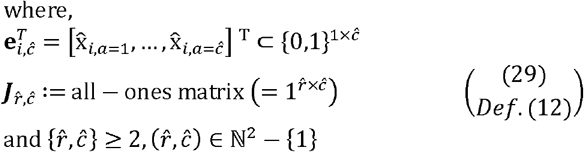

The contributions, if any, for each member of a ligand-interacting macromolecular species can be summarized by the column sum of the occupancy matrix,

#### Lemma 2 (**L2; without proof**)

The distribution of 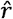-ligand-interacting macromolecules across *ĉ*-conformational states, for an ensemble distribution, is,

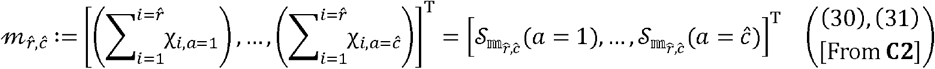

where,

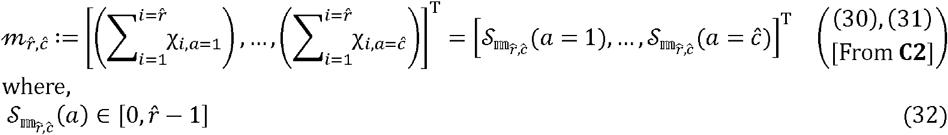

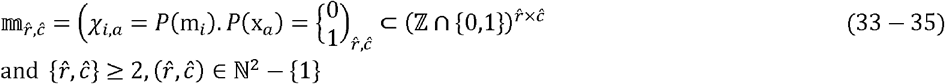

#### Corollary 3 (**C3**; **without proof**)

The p1-norm of 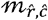, for an ensemble distribution, is equal to the number of ligand-interacting identical macromolecules that constitutes it,

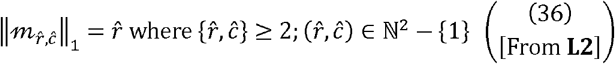

The occupancy matrix 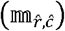, for an ensemble distribution, is parameterized as a multinomial distribution with the probability 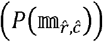,

#### Theorem 1

(**T1;** Section 6, **Eqs. (21-27)**):

The probability of the distribution matrix 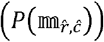, for an ensemble distribution, is the multinomial distribution of 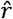-ligand-interacting macromolecules across *ĉ* - conformational states,

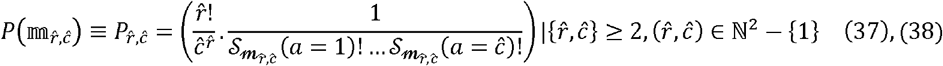

We now summarize the properties of the occupancy matrix 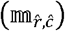, with formal definitions/derivations, for an ensemble distribution; (Step 1, **Figure 1**),

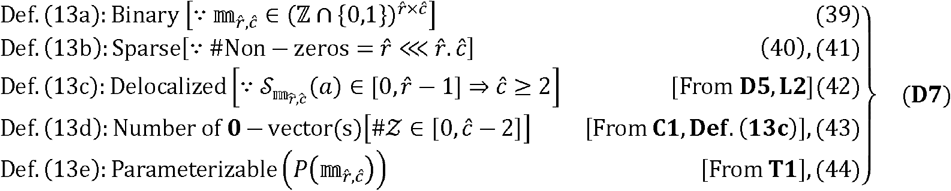

**Figure 1:**
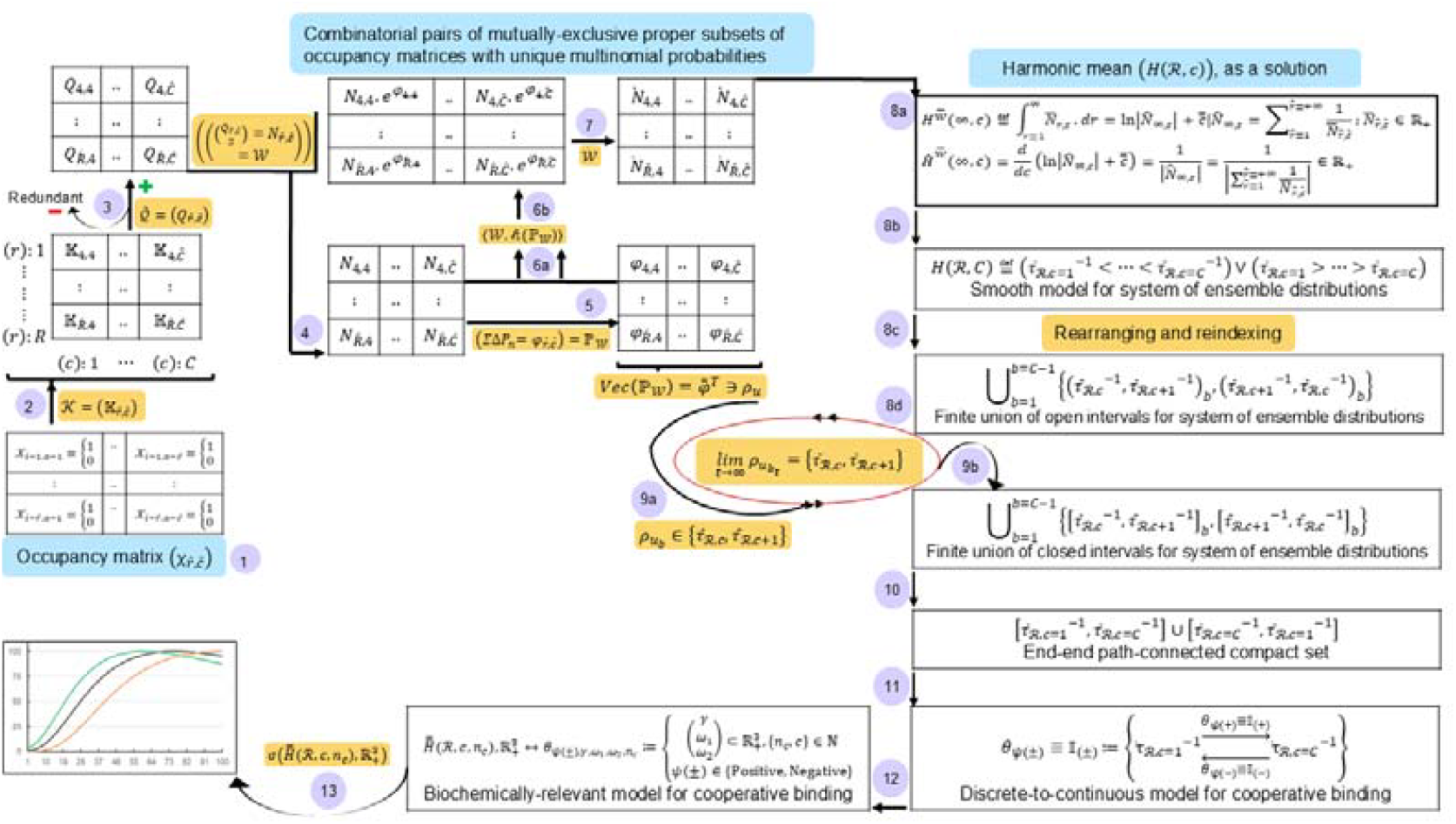
Algorithm to construct the discrete-to-continuous mathematical model for a system of ensemble distributions of ligand-interacting macromolecular species across milieux-dependent conformational states: The premise, of our model, are partitioned by a multinomial probability-based hyperparameter into mutually exclusive proper subsets of occupancy matrices (sparse, binary, strictly delocalized), for an ensemble distribution (species of ligand-interacting identical macromolecules, milieux-dependent conformational states), with identical multinomial probabilities (Steps 1-3). Since each subset is canonical with a constituent occupancy matrix, it is characterized by a unique multinomial probability. The inner product of the matrices (number of combinatorial pairs of subsets, expression for the summed transitional probabilities) is the differentiable positive real-valued matrix for the system of ensemble distributions (Steps 4-7). Whilst the harmonic mean is presented as a possible solution for the system of ensemble distributions, the row-wise definite integral for each column is computed (Step 8). An algebraic representation, for the latter, is the finite union of open intervals (contiguous, strictly monotone) which in tandem with the set of interval-specific and bounded transitional probabilities results in a piecewise smooth curve (path-connected-, closed-, compact-set) (Steps 9-11). The resulting- and modified (coefficients, exponents)-curve is bijective with the sigmoid function and can be assessed for biochemical relevance (Steps 12 and 13). A**bbreviations**: system-level details (indices, cardinality) for the system of ensemble distributions;, sigmoid-function mapped and parameterized piecewise smooth curve with regular- and smooth-closed intervals as milieux-dependent conformational states;

### 2.3 Finite set of occupancy matrices for an ensemble distribution

The constraints outlined in **D7** suggest that 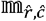 is amenable to substitution-without-replacement (factorial) which can be deployed to generate an ensemble of occupancy matrices each of which is parameterized with a multinomial probability (**T1**). An analysis of the probabilities, post hoc, suggests the following distribution,

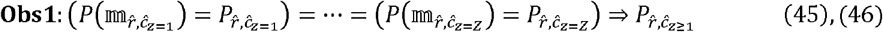

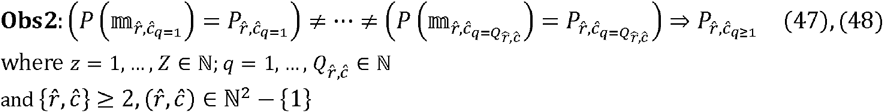

We present a partitioning schema for the set of all possible probabilities, for an ensemble distribution, by combining **Eqs. (45-48)**.

#### Theorem 2 (**T2**; **without proof**)

For an ensemble distribution, the set of multinomial probabilities can be partitioned into *q*-indexed 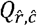-subsets 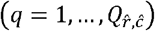 of *Z*_*q*_-membered identical multinomial probabilities,

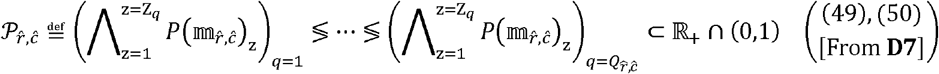

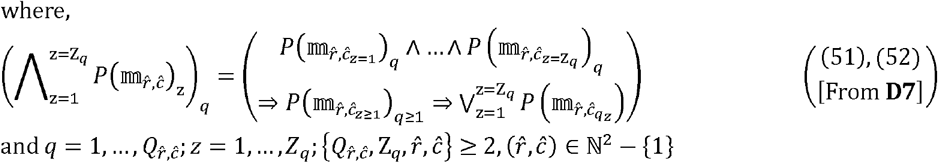

The partitioning of the occupancy matrices, for an ensemble distribution, can be inferred,

#### Corollary 4 (**C4; without proof**)

For an ensemble distribution, the set of occupancy matrices can be partitioned into *q*-indexed 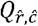-subsets of *Z*_*q*_-membered occupancy matrices,

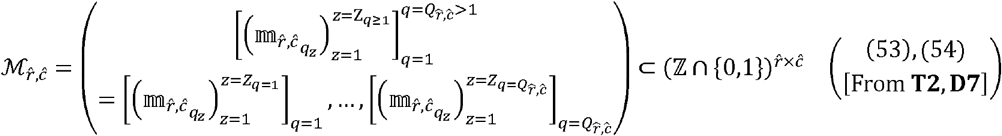

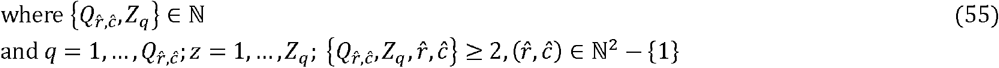

#### Corollary 5 (**C5.1-C5.4; without proof**)

For an ensemble distribution, there exists a set of -indexed occupancy matrices 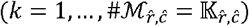,

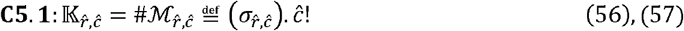

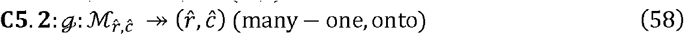

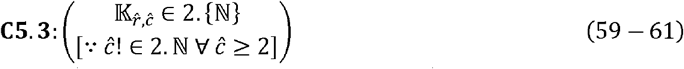

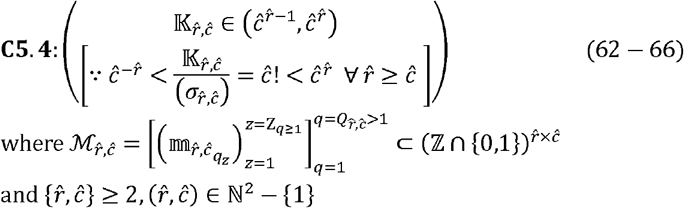

Paraphrasing the distribution of multinomial probabilities outlined in **T2** in terms of *k*-indexed 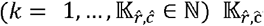-occupancy matrices (**C4**), for an ensemble distribution; (Step 2; **Figure 1**),

#### Corollary 6 (**C6; without proof**)

For an ensemble distribution, the set of -indexed 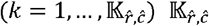 -multinomial probabilities 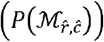 can be partitioned into *q*-indexed 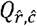 -subsets of *Z*_*q*_-membered multinomial probabilities,

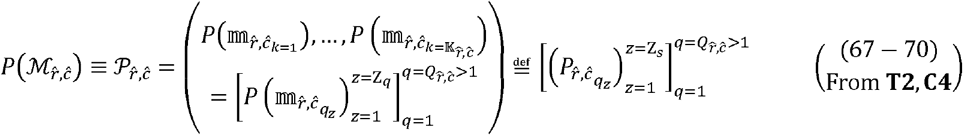

where,

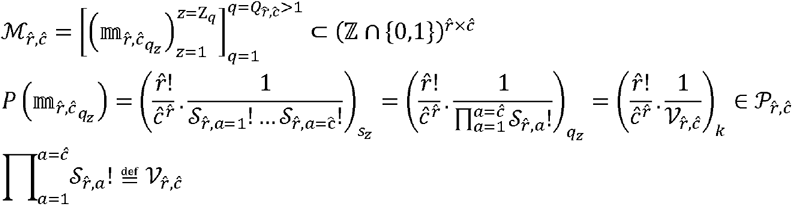

With 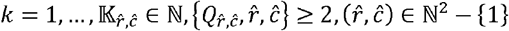

For greater clarity on the aforementioned schema, we present results for a subsequence of probabilities and complement the same with solved numerical examples

#### Corollary 7 **(C7; without proof)**

For an ensemble distribution, the multinomial probabilities for *q*-indexed 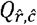 -terms 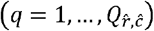, can be ordered as strictly increasing- or decreasing-subsequences 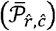 of -indexed _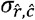_-unequal terms 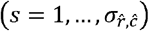,

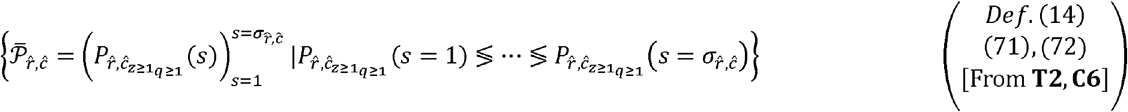

where,

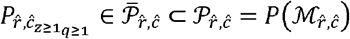

and

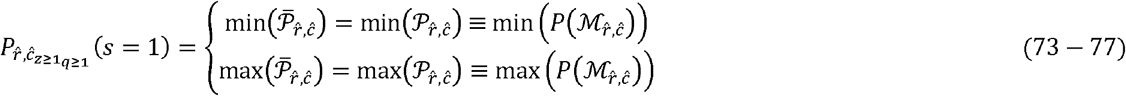

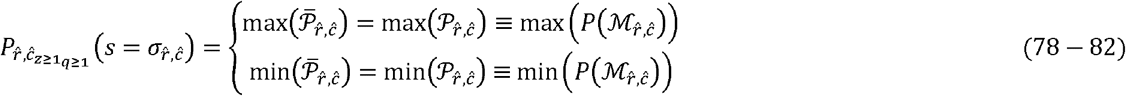

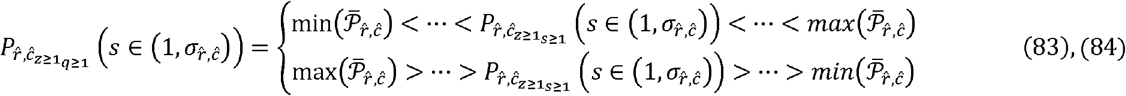

with 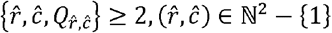

We present solved examples for greater clarity of the definitions and results in this section,

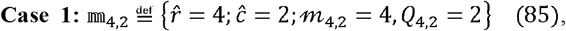

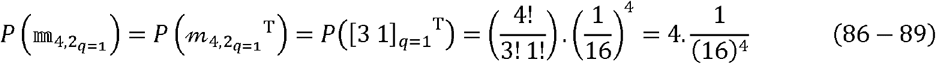

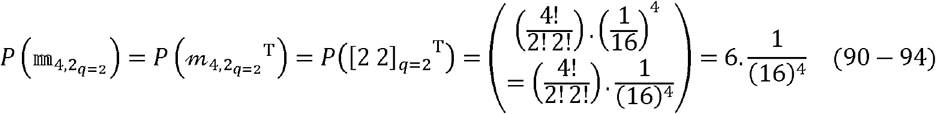

From Eqs. (89) and (94)

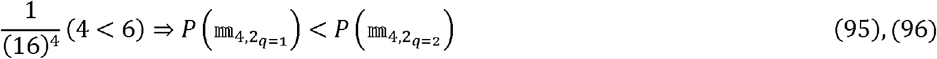

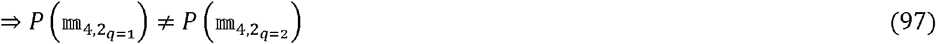

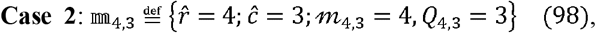

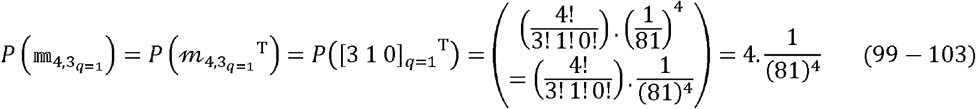

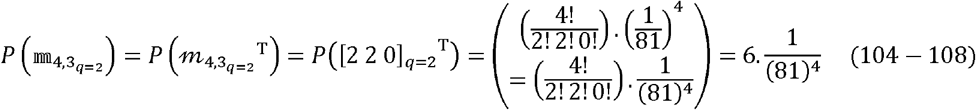

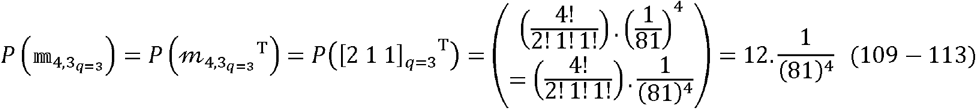

From Eqs. (103),(108) and (113)

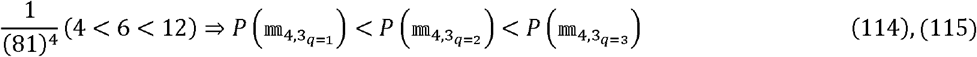

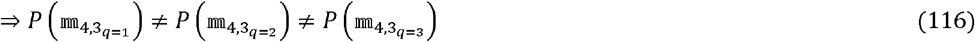

Whilst the results, vide supra (**T2, C4** − **C7**), were observational, we define a probability-based hyperparameter 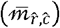 which is based on the product-of-column-sums to ensure mathematical rigor,

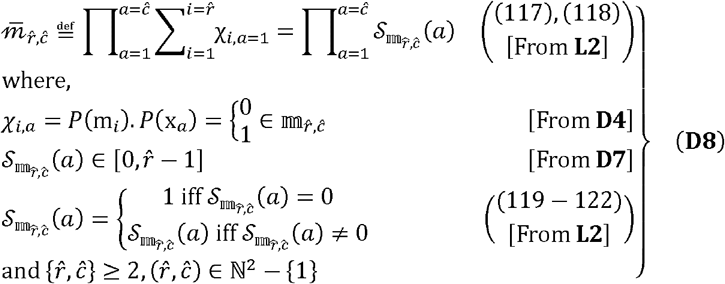

We now deploy 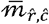 and partition the set of unequal occupancy matrices into the following subsets in accordance with **T2** and **C4-C7**, for an ensemble distribution,

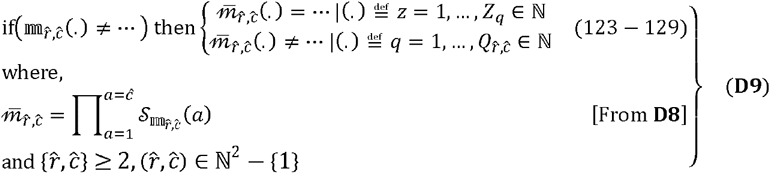

We summarize the aforementioned results, for an ensemble distribution, with the assertion that the probability-based hyperparameter 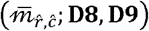 will partition the set of 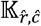-occupancy matrices on the basis of their multinomial probabilities (identical, unique) (**T2, C4-C7**).

### 2.4 Properties of 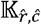-occupancy matrices for an ensemble distribution

Whilst **C5** discussed 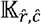 numerically (cardinality, composition), here, we evaluate the same qualitatively, i.e., in terms of the properties of the occupancy matrices that comprise 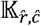.

#### Lemma 3.1 (L3.1; **without proof**)

For 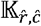 -occupancy matrices, of an ensemble distribution, the -*q*^*th*^ subset is proper,

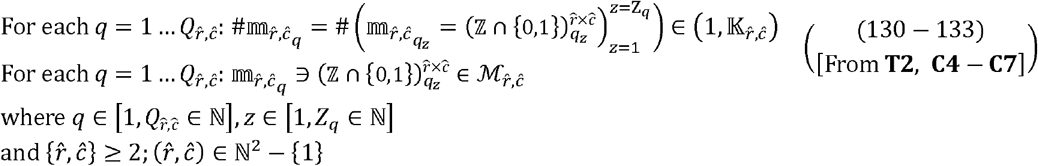

#### Lemma 3.2 (L3.2; **without proof**)

For an ensemble distribution, with 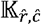-occupancy matrices, the *q*-indexed 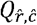-proper subsets 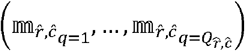 are mutually exclusive,

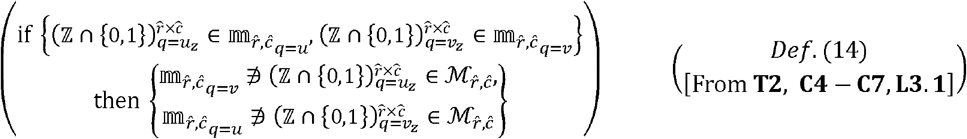

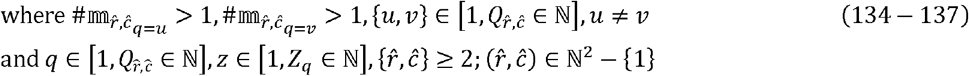

#### Lemma 3.3 (**L3.3**; **without proof**)

For an ensemble distribution, with 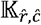-occupancy matrices, the *q*-indexed 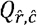-proper and mutually exclusive subsets 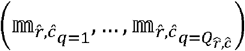 may be regarded as canonical,

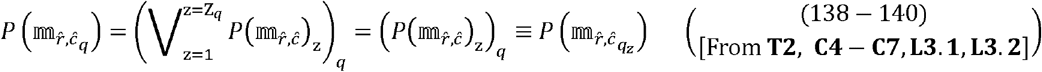

where 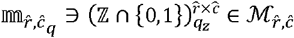

and 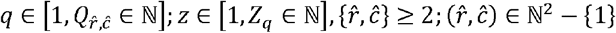

#### Lemma 3.4 (**L3.4**; **without proof**)

The multinomial probabilities associated with the *q*^th^-subset of 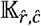-occupancy matrices, for an ensemble distribution, is unique,

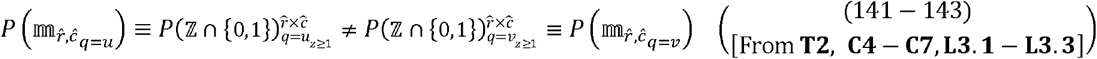

where,

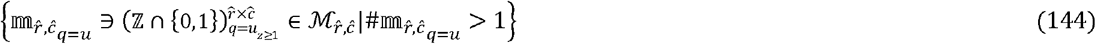

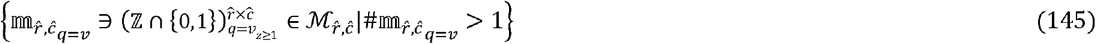

and 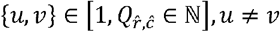

with 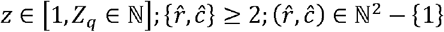

We now define and derive a revised index 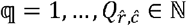 for an ensemble distribution with 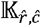-occupancy matrices which will be utilized, hereafter, without any loss of generality, for reduced ambiguity and improved clarity.

#### Theorem 3 (**T3**; **without proof**)

The index 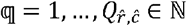, can be utilized to represent 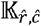-occupancy matrices, for an ensemble distribution, without any loss of generality.

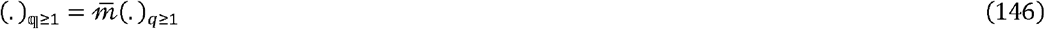

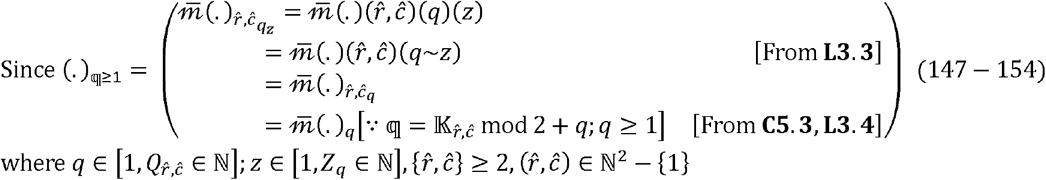

The 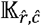-occupancy matrices, for an ensemble distribution, can be partitioned by a probability-based hyperparameter 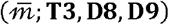 into q-indexed mutually exclusive proper subsets each of which is canonical with any constituent occupancy matrix and can therefore, be characterized by a unique multinomial probability. Formally,

#### Corollary 8 (**C8**; **without proof**)

The q^*th*^-mutually exclusive and proper subset, for an ensemble distribution, is canonical and is characterized by a unique multinomial probability,

m_q_ is proper subset:

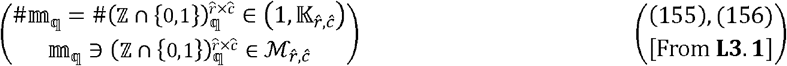

m_q_ is a mutually exclusive subset:

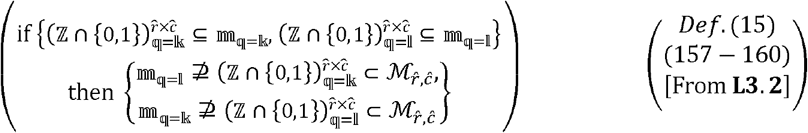

m_q_ is canonical:

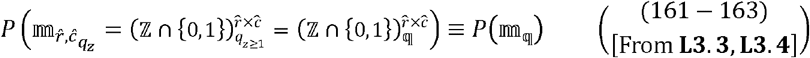

m_q_ is uniquely characterized:

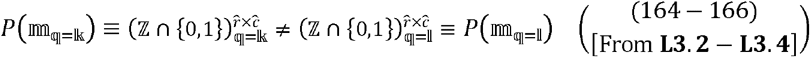

where,

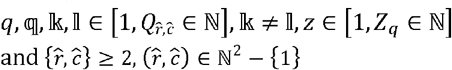

### 2.5 Thermodynamic considerations for a discrete-to-continuous model of an ensemble distribution

The role of thermodynamic fluctuations in the genesis of cooperative binding is an important consideration and cannot be understated. For a statistical ensemble that comprises -particles this is likely to include the intra (macromolecules, small molecule modifiers)- and extra (physicochemical)-cellular microenvironments under standard (temperature, pressure, volume)-conditions,

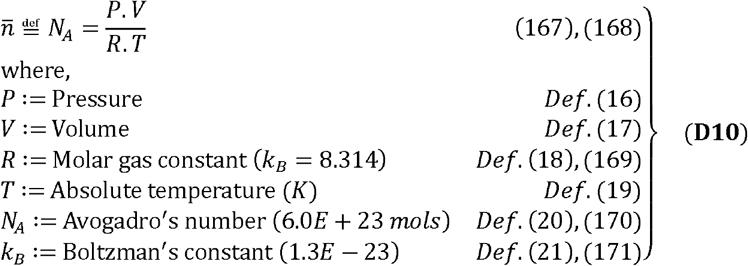

The vibrational frequency of the participating atoms may be deployed as an index of atomic fluctuations and thence complex function [13-15],

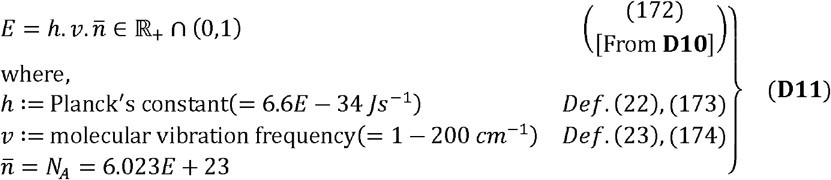

We now derive an expression to compute the potential energy in a milieu of 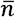-total particles,

#### Lemma 4

(**L4;** Section 6, **Eqs. (28-39)**):

The potential energy for a system with 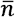-total particles,

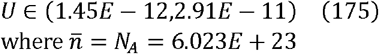

We revise our definition, for a conformational state, and describe the same as non-colligative, “milieu-dependent” and parameterizable with a unique probability,

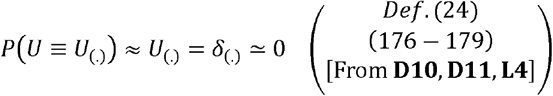

On applying these assertions and results to our ensemble distribution of q-indexed 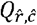-mutually exclusive proper subsets with unique multinomial probabilities, we get,

#### Lemma 5 (**L5; without proof**)

The theoretical estimate for the probability of *U*_q_ is the unique multinomial probability associated with 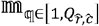,

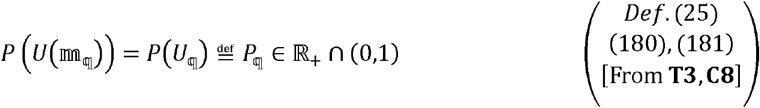

where 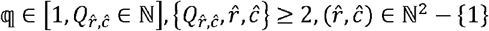

Extending **L5** for an ensemble distribution 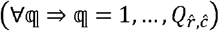

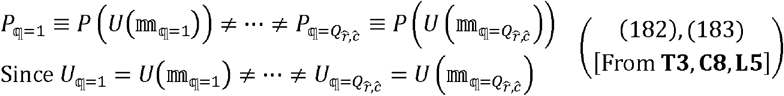

In order to retain the thermodynamic relevance of our discrete-to-continuous mathematical model, for an ensemble distribution, the potential energies and the corresponding multinomial probabilities for any pair of mutually exclusive proper subsets is formulated and expressed (**T3, C8, L5**).

#### Corollary 9

(**C9;** Section 6, **Eqs. (40-61)**):

For any distinct k,l-pair of q-indexed mutually exclusive proper subsets 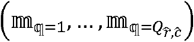 the corresponding potential energies and thereby its multinomial probabilities are unique,

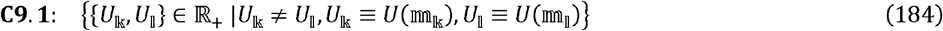

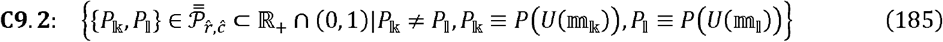

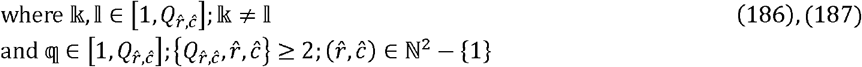

## 3 Results

In this section we present expressions, estimates, bounds, metrics and indices for ensemble distributions of ligand-interacting macromolecular species across milieux-dependent conformational states.

### 3.1 Discrete model for ensemble distribution(s) of a ligand-interacting macromolecular species across milieux-dependent conformational states

We have modeled, parameterized and partitioned the set of occupancy matrices 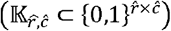, for an ensemble distribution, into 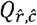-mutually exclusive proper subsets 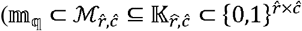 whilst ensuring the thermodynamic relevance of the same (**D8, D9, C8, C9**). Here, we combine these assertions and extend the same to a complete system (subsections: 3.1.1-3.1.3).

#### 3.1.1 Bounds for the set of mutually exclusive proper subsets for an ensemble distribution

In keeping with our premise to model macromolecular plasticity, our modeling effort requires utilizing combinatorial pairs of 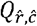-mutually exclusive proper subsets 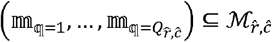 to model probable scenarios; (Step 3; **Figure 1**),

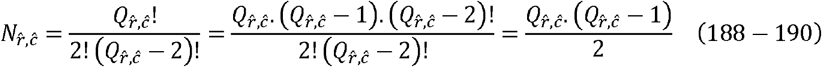

where,

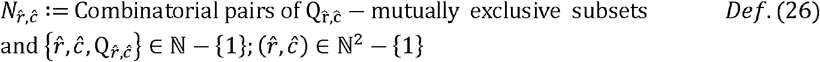

We now compute lower bounds for 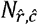 and 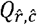 by solving **Eq. (190)**,

##### Corollary 10

(**C10;** Section 6, **Eqs. (62-76)**):

For an ensemble distribution with 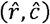, the lower bound for 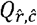,-mutually exclusive proper subsets is dependent on 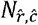- combinatorial pairs,

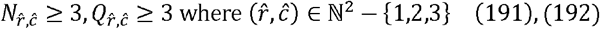

Since each subset is canonical with a constituent occupancy matrix and is therefore, characterized by a unique multinomial probability, the finite difference between the unique multinomial probabilities for any pair of 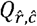-mutually exclusive proper subsets is defined as a transitional probability (***Def. (27)***),

##### Lemma 6

(**L6;** Section 6, **Eqs. (77-90)**):

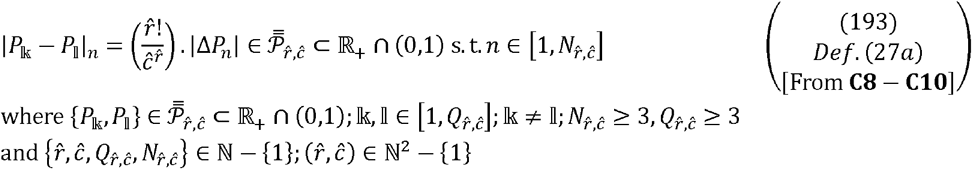

#### 3.1.2 Cardinality for the set of mutually exclusive proper subsets for an ensemble distribution

Computing the number (cardinality) of 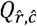-mutually exclusive proper subsets, for an ensemble distribution with 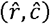, is a fundamental step in developing our discrete-to-continuous mathematical model. We extend the previously defined generic-properties and -constraints outlined in **D7**, to include the following specific constraints whence evaluating and enumerating the cardinality for the same,

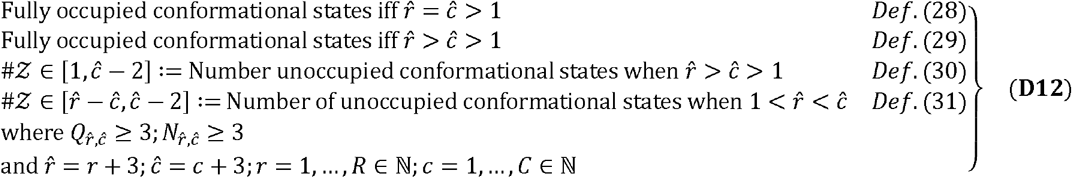

Combining **D7** and **D12** we get,

##### Lemma 7 (**L7, without proof)**

The cardinality for 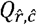--mutually exclusive proper subsets, for an ensemble distribution with 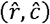, is dependent on the choice of indices for 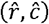,

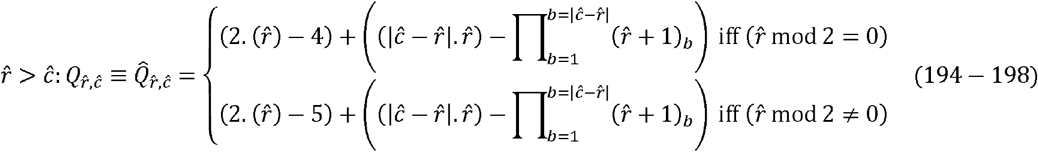

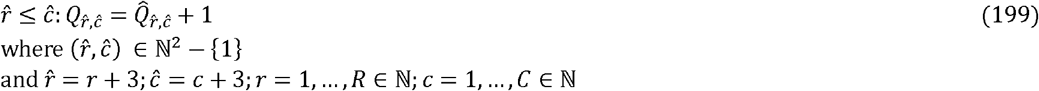

where 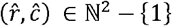
and 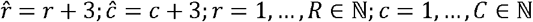

#### 3.1.3 Discrete model for the system of ensemble distributions

We utilize the aforementioned results to construct a system of ensemble distributions of ligand-interacting macromolecular species across milieux-dependent conformational states which we formally define as the *R* × *C*-strictly positive matrix where each entry is indexed by 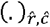. We have seen that the choice of indices 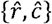 is an important step in developing our discrete-to-continuous model, which, in turn, is contingent upon the constraints (**D7, D12**) and lower bounds (**C10, L7**). Our revised choice of indices, is computed with,

##### Lemma 8

(**L8**; Section 6, **Eqs. (91-114)**):

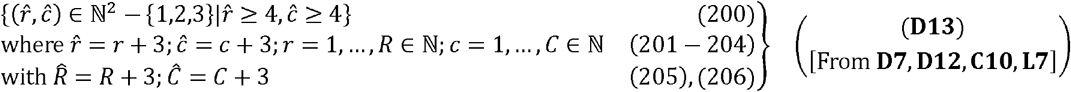

We now define and populate the appropriate system-level matrices,

##### Lemma 9 (**L9; without proof**)

The system-level finite matrices {ℕ, ℝ_+_}^*R*×*C*^ can be expressed in terms of computed indices 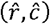,

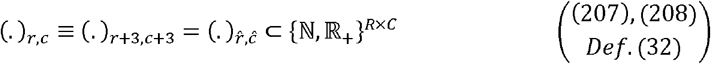

where

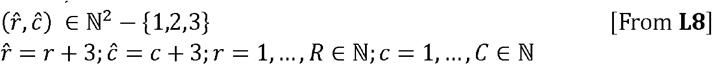

and,

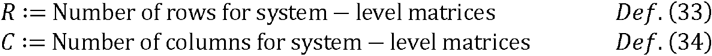

The system-level matrices 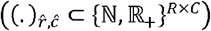 that are pertinent to our discrete-to-continuous model of ensemble distributions of ligand-interacting macromolecular species across milieu-dependent conformational states are defined and populated,

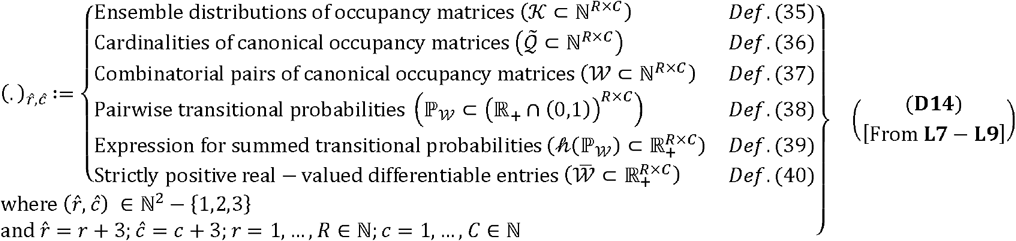

### 3.2 Discrete-to-smooth transformation for the system of ensemble distributions

The development of our discrete-to-continuous mathematical model is dependent on a transformation schema wherein the discrete systems described, vide supra, are transformed into one that allows for smooth descriptors and analyses (subsections: 3.2.1-3.2.3).

#### 3.2.1 Combinatorial pairs of mutually exclusive proper subsets, of ensemble-specific data, for the system of ensemble distributions

We had previously discussed the derivation and computation for 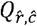 with a quadratic expression for different values of 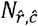 and 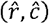 (**L7-L9**). Here, we extend the same to all ensemble distributions with the system-level *R* × *C*-matrix of natural numbers (*W*) (**D14**); (Step 4; **Figure 1**),

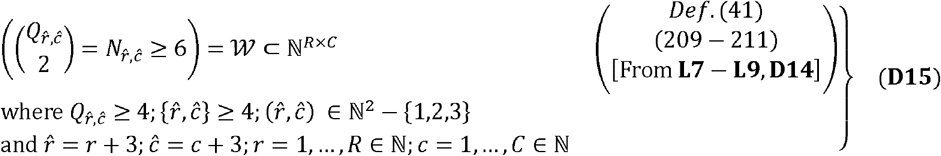

#### 3.2.2 Transitional probabilities, between mutually exclusive proper subsets of ensemble-specific data, for the system of ensemble distributions

Since *W* represents the cardinality of combinatorial pairs of canonical occupancy matrices, the finite differences or transitional probabilities between all 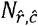-pairs can be summed and will constitute the system-level *R* × *C*-matrix of strictly positive real-valued numbers between 0,1 (ℙ _*W*_ ⊂ (ℝ _+_ ∩ (0,1)) ^*R* × *C*^) (**D14**); (Step 5; **Figure 1)**,

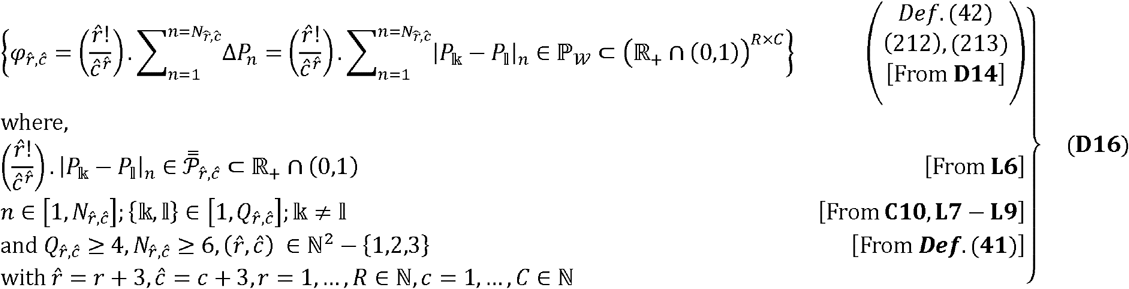

#### 3.2.3 Discrete-to-continuous transformation, of ensemble-specific data, for the system of ensemble distributions

We seek a strictly positive real-valued representation of 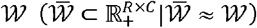, for the system of ensemble distributions, which will ensure differentiability, specificity and consistency of the chosen coefficient.

##### Lemma 10 (**L10**; **without proof):**

For every 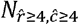, we can identify a unique positive real-valued coefficient 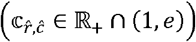 which is consistent and annotatable.

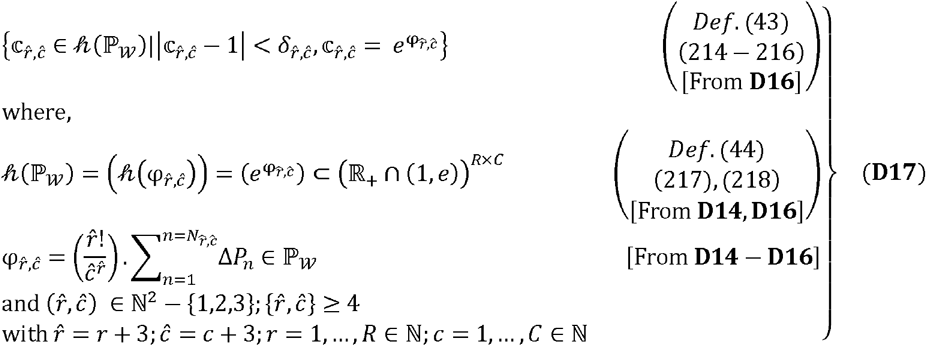

We can immediately appreciate the following results for 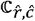

Corollaries 11.1-11.3 (**C11.1-C11.3**; **without proof**): The coefficient 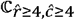 for 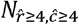 is a unique positive real-valued number 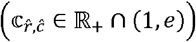 which is variable, consistent and annotatable.

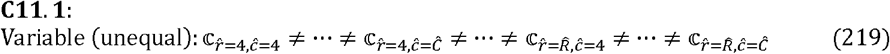

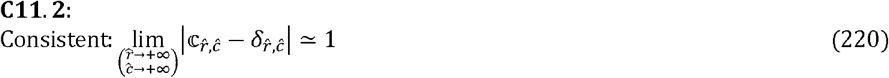

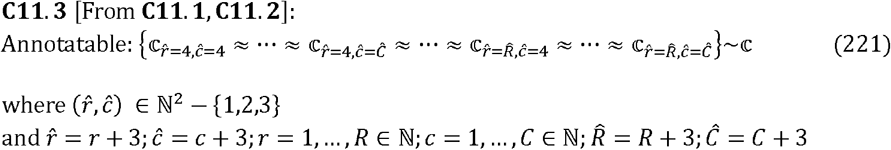

Rewriting these results as a linear map and expressing the same in matrix notation in accordance with **D14**; (Step 6; **Figure 1**),

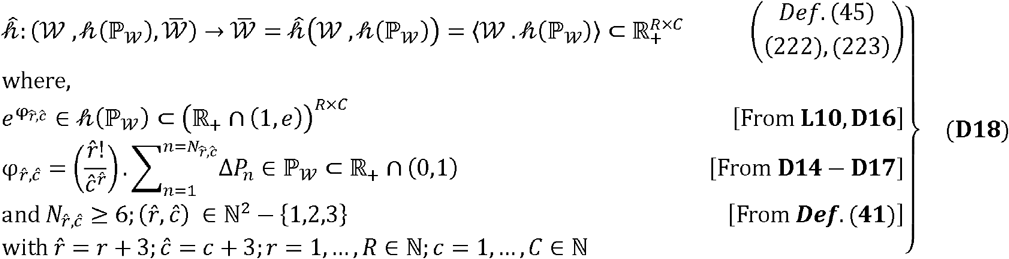

Expanding 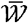 for each 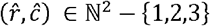; (Step 7; **Figure 1**),

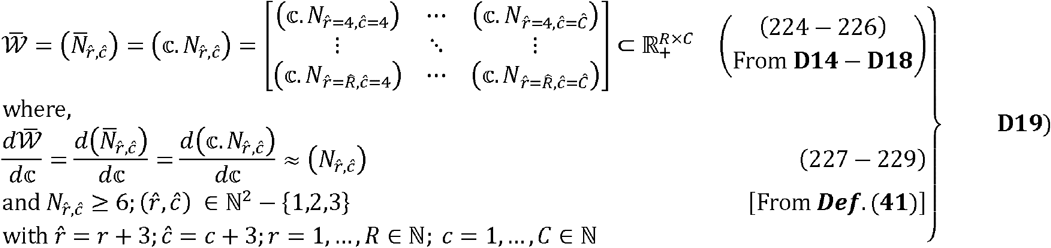

### 3.3 Systematic row-wise sampling per column for the system of ensemble distributions

We have discussed the relevance of the milieu both, external (physicochemical) and internal (biochemical) in the genesis of cooperative binding, for an ensemble distribution, and thence extended the same to all such distributions via the system-level matrices (**L9, D14**). However, in order to evaluate the role of our model in the progression of cooperative binding we will need to demonstrate that our representation is piecewise smooth in order to recapitulate cooperative binding. We will accomplish this by systematically sampling the ligand-interacting macromolecular species, row-wise for a large *R* ∈ ℕ, i.e., any *c*^*th*^ -column,

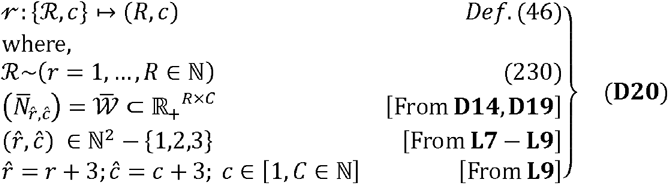

We seek a function to model, characterize, analyse and thence map our system of ensemble distributions to known empirical models of the same (subsections: 3.3.1-3.3.3).

#### 3.3.1 The Harmonic mean, as a solution, for the system of ensemble distributions

We deploy the Harmonic mean systematically for different cardinalities/species per column, for a possible solution, and then extend the same across all columns for a generic solution.

##### Theorem 4

(**T4;** Section 6, **Eqs. (115-125)**):

The Harmonic mean, for the *c*^*th*^ -column of 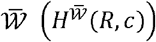 is a solution of non-zero 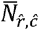-real values that comprise the same,

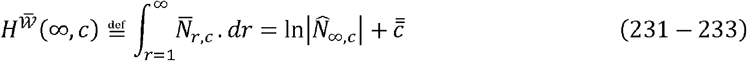

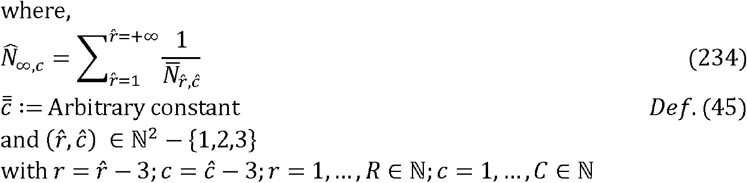

##### Corollary 12 (**C12**; **without proof):**

The differential equation, for the *c*^*th*^ -column, of the differential system 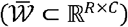, is a non-zero positive real number,

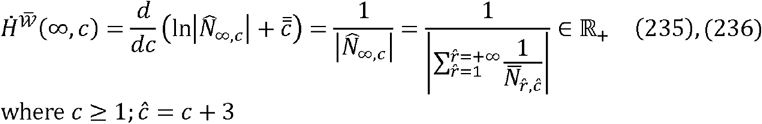

From **T4** and **C12** we are able to assert that the *c*^*th*^ -column is smooth, a result that is extendible to all *C*-columns of 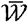 (Step 8; **Figure 1**).

#### 3.3.2 Characterizing the system of ensemble distributions

In order to explicitly characterize each conformational state, for any given *R* ∈ ℕ, we define and deploy the row-wise definite integral,

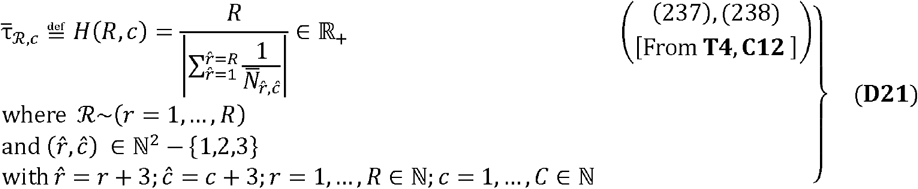

Extending these results to all *C*-columns we observe that *H* (*R, C*) is a strictly monotone (decreasing, increasing) sequence (Step 8a; **Figure 1**),

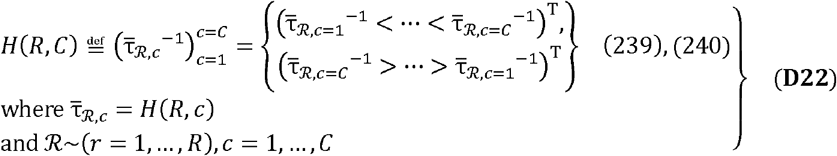

#### 3.3.3 Interval representation for the system of ensemble distributions

Since *H* (*R, C*) comprises points that are monotone, continuous, and strictly increasing or decreasing, we can index any contiguous pair and reannotate this as a real-valued and bounded open interval (I_*b*_),

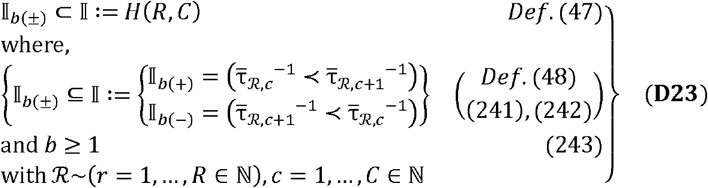

We now express the system of ensemble distributions *H* (*R, C*) as the set of contiguous and monotone open intervals; (Step 8b, **Figure 1**).

##### Corollary 13 (**C13**; **without proof**)

*H* (*R, C*) can be expressed as the finite union of *b*-indexed *C* − 1-contiguous and monotone open intervals,

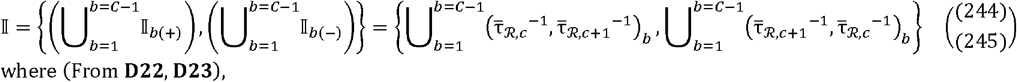

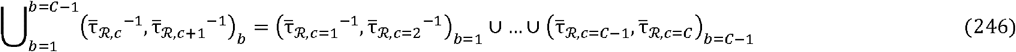

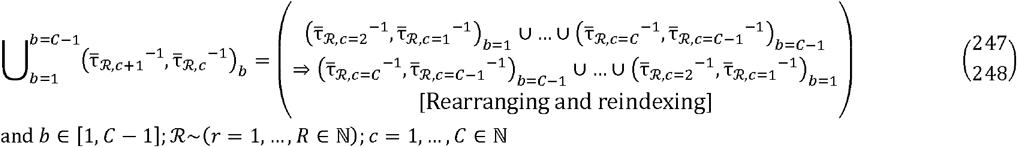

### 3.4 Transition probability-based path-connectedness for contiguous and monotone open intervals

The final step in our discrete-to-continuous model is to establish *H* (*R, C*) as a compact set. In order to unequivocally prove this, we will need to demonstrate that each contiguous open interval (pair of unequal “delay”-metrics) is path-connected and therefore, closed. We will deploy the previously defined matrix of transitional-probabilities (ℙ_*W*_) to establish continuity (subsections: 3.4.1-3.4.3)

#### 3.4.1 Bounds for the system of transitional probabilities

In order to fully utilize the transitional probabilities ℙ_*W*_ ⊂ ∈_+_ ⋂ (0,1), we will need to define the specific bounds from where to make this selection.

##### Lemma 11

(**L11;** Section 6, **Eqs. (126-135)**):

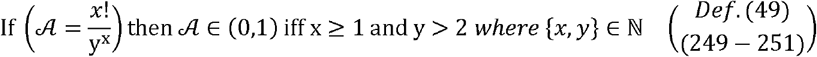

##### Theorem 5

(**T5;** Section 6, **Eqs. (136-159))**:

For the ordered-pair 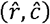,

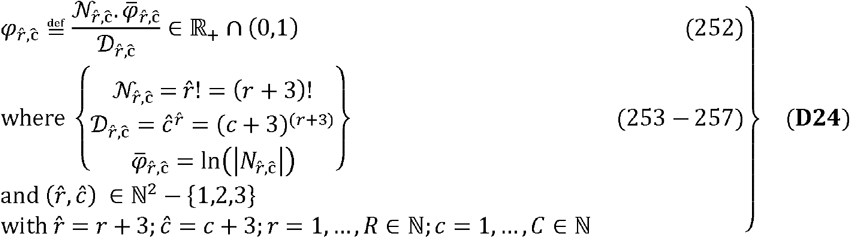

##### Corollary 14

(**C14;** Section 6, **Eqs. (160-172)**):

The lower bound for the probability mass function of a complete system ℙ_*W*_ is,

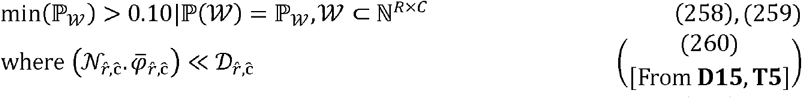

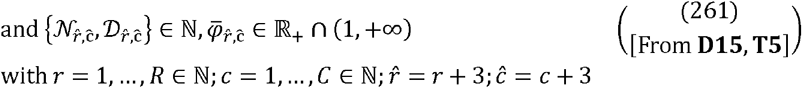

We are now in a position to extend these results for a large number 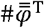 which we define as the cardinality for the vector of all pooled q-indexed 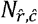-combinatorial pairs of transitional probabilities; (Step 9a, **Figure 1**),

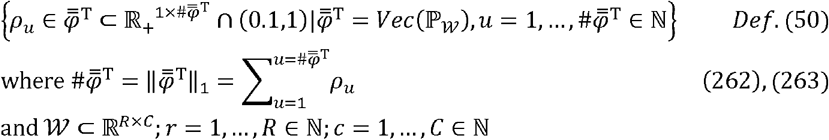

##### Theorem 6

(**T6;** Section 6, **Eqs. (173-206)**):

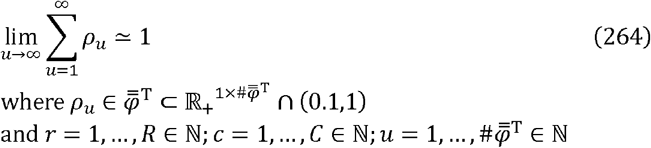

#### 3.4.2 Interval-specific and transitional probability-based sequence(s) of terms

We have, vide supra, defined the set of transitional probabilities and established its boundedness (**T5, T6, C14**). We now utilize the same to establish continuity for each monotonic and contiguous open interval of (*H* (*R, C*)).

Let us compute and annotate any transitional probability as interval-specific 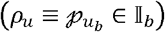,

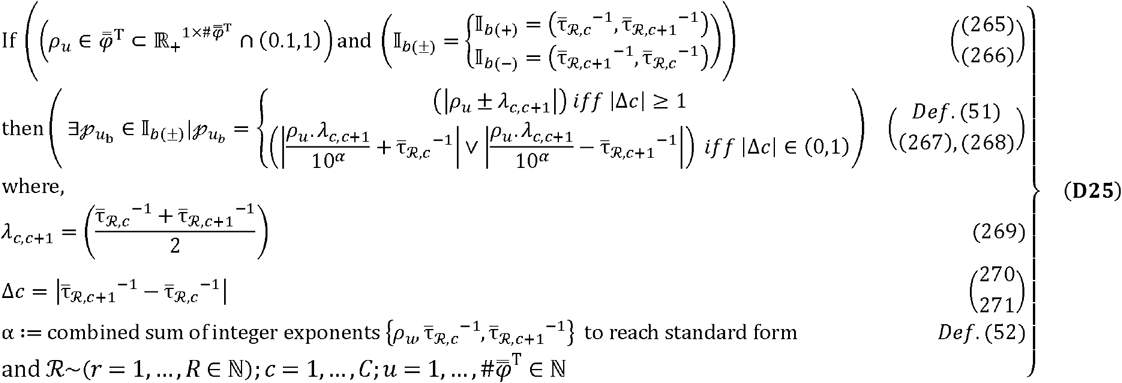

We now purport the existence of interval-specific and transitional probability-based infinite sequences for any open interval with the following results; (Step 9b; **Figure 1**),

##### Theorem 7

(**T7;** Section 6, **Eqs. (207-231)**):

For any 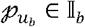, we have a *t*-indexed sequence,

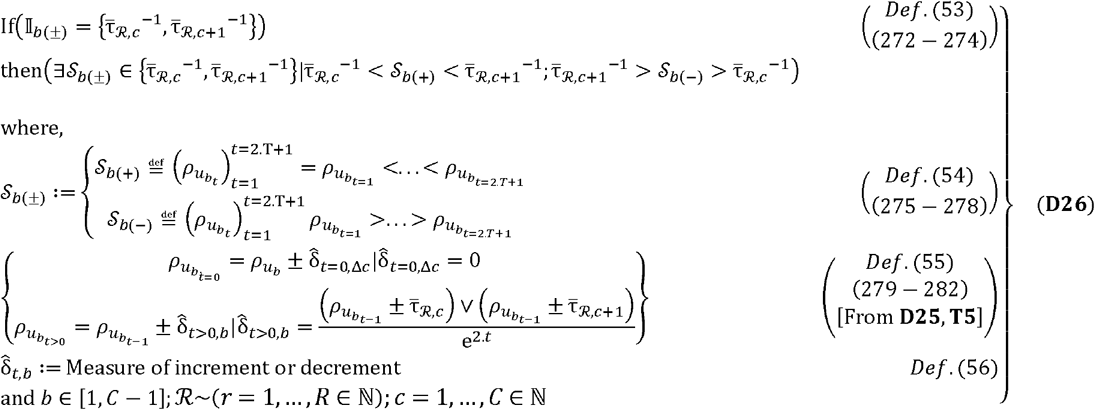

We can extend **T7** to include infinitely many terms with,

##### Corollary 15 (**C15; without proof)**

For any 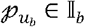 we have 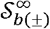, the superset of -indexed sequences,

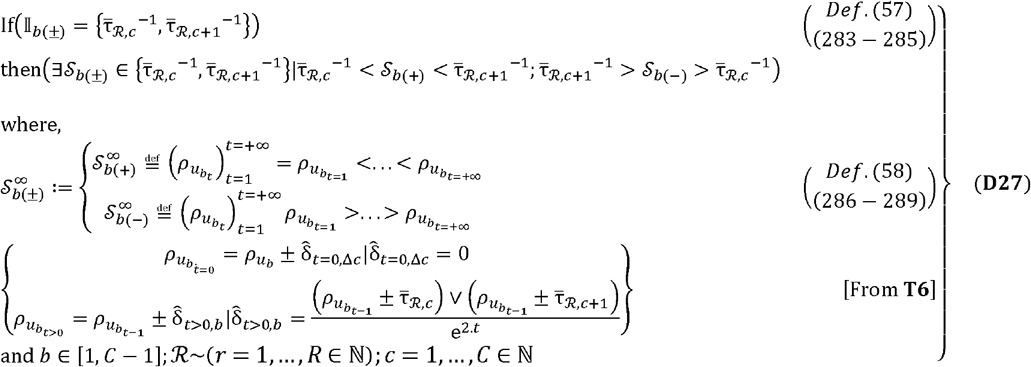

We can assert, from **T6, D25, T7** and **C15**, that with the exception of 2. *T* + 1 terms, there are infinitely many terms for any interval-specific transitional probability (**D25**-**D27**).

#### 3.4.3 Convergence of transitional probability-derived interval-specific terms

We now utilize any interval-specific transitional probability and the sequence(s) of infinite terms that are associated with each to establish path-connectedness and convergence (Step 9a; **Figure 1**).

##### Theorem 8 (**T8; without proof)**

The t-indexed infinite sequence 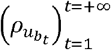 of interval-specific transitional probabilities converges,

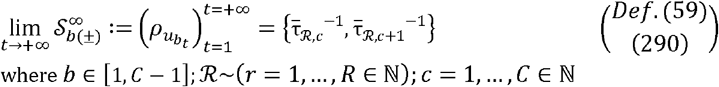

##### Corollary 16 (**C16; without proof**)

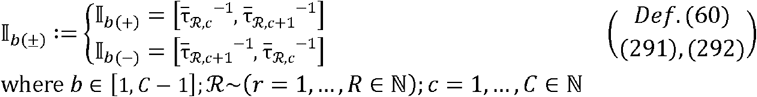

##### Corollary 17 (**C17; without proof**)

I_*b*(+)_ and I_*b*(−)_ are path-connected and therefore, compact [From **C16** and **HB theorem**].

Combining, rearranging and reindexing these steps we can immediately state that 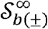 converges for all *C* − 1 intervals; (Steps 9b and 8d; **Figure 1**).

##### Corollary 18 (**C18; without proof**)

Each *b*-indexed *C* − 1-interval (I_*b*=1_, …, I_*b*=*C*−1_) is path-connected and compact.

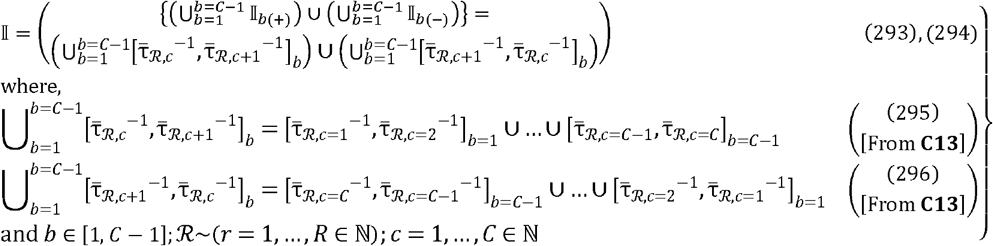

##### Corollary 19 (**C19; without proof**)

The set formed by the finite union of all compact intervals, of *C*-conformational states is an end-end closed interval and therefore, compact; (Step 10, **Figure 1**)

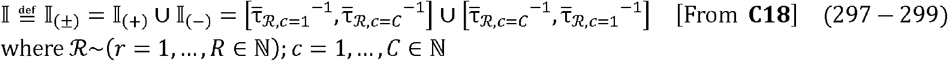

##### Corollary 20 (**C20; without proof**)

The end-end compact set can be partitioned into well-defined compact intervals and is therefore, piecewise smooth [From **C15** – **C21**].

## 4 Discussion

Whilst the presented assertions, reasoning and analyses are sound and rigorous, the biochemical relevance of our discrete-to-continuous model remains to be ascertained. Here, we extend our results and discuss potential applications of our model, in the presence of empirical data, whence representing complex biochemical systems.

### 4.1 Incorporating system specificity and response into our discrete-to-continuous model

Our discrete-to-continuous model is a compact set and can be described as a curve which is piecewise smooth (**C16-C20**). However, in order to model the kinetics of complex biochemical systems, parameterizing the same is necessary to ensure context and thence biochemical relevance (subsections: 4.1.1-4.1.3).

Since our model is a piecewise smooth curve, it may be apt at modelling both, positive- and negative-cooperative binding (Step 11; **Figure 1**). Paraphrasing this,

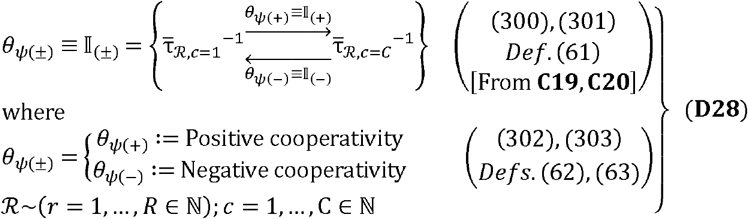

#### 4.1.1 Parameterizing abscissa-values for the sequential assessment of a complex biochemical system

Whilst the magnitude of a biochemical response varies in proportion to the incident stimulus, the response from an underlying biochemical systems is usually temporal and will decay. For example, the restoration of precession frequency in proton-NMR (nuclear magnetic resonance) after a radio-frequency (RF)-stimulus, time-of-flight (TOF) of samples in matrix-assisted laser desorption ionization (MALDI)-based mass spectrometry (MS, MS/MS) and radioisotope studies (labeled blotting, immunoassays) are computed parameters where signal data is collated from area-under-curve (AUC)-measurements of time-series data [63-66]. The abscissa, additionally, is also utilized to represent the underlying biochemical heterogeneity, for the aforementioned analyses, as analyte concentrations (NMR, water, lipid, bone; radioisotope studies, metabolites of the glycolytic pathway) and absorbance maxima (*λ*_*max*_; UV-Vis) for complex biochemical solutions (DNA/RNA, proteins, cytochrome c, hemoglobins, NAD- and FAD-based such as false discovery rate (FDR) which is *min* (*FDR*_1_,_…_,) from a multiplicity of tests or repeated coenzymes) [65, 66]. The advent of big data has also redefined the AUC to compute hyperparameters sampling in order to retain biochemical relevance [67-70]. This peak or the lack thereof of one, is a quantitative measure of the total number of molecules that exhibit a certain characteristic or state.

Here, we introduce the set of natural numbers, into the Harmonic mean, in an effort to render our discrete-to-continuous model amenable to incremental biochemically- and physiologically-relevant factors,

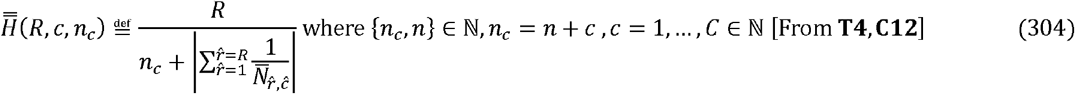

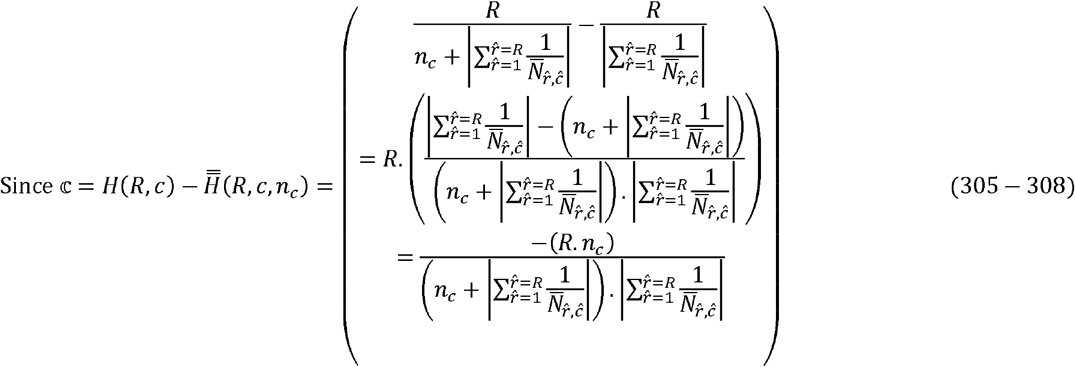

#### 4.1.2 Parameterizing ordinate-values to model response for a complex biochemical system

The signal strength, whence parameterized independently from the AUC is either a peak or plateau and is defined as the maximum strength/magnitude/intensity of response to an incident stimulus. Conversely, comparative analyses of the same can offer improved resolution, reduced ambiguity and a filtering criteria. For example, the molar extinction coefficient (*ε*), for mixtures of biological macromolecules preclude wavelengths (*λ*_*max*_) with poor resolving power for any number of candidate macromolecules with complex intramolecular geometries can be utilized under different experimental conditions to [65-68]. Perfusion studies, drug bioavailability assays for adverse drug reaction monitoring and dosing studies, peak gastric output, reaction velocities for enzyme-based conversions, plasma glucose levels in tandem with oral glucose tolerance tests (OGTTs) and oxygen saturation levels are examples of system-specific assays that deploy signal intensity as independent parameters to assess the physiological- and biochemical-function under study [71-73].

Our discrete-to-continuous mathematical model, for a system of ensemble distributions, is a piecewise smooth curve which can be numerically modified with strictly positive real-valued coefficients to model peak signal intensity as an independent determinant of biochemical function; (Step11, **Figure 1**),

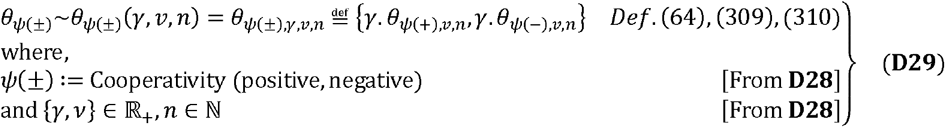

#### 4.1.3 Sigmoid function as the activation function for our discrete-to-continuous model

The modifications for abscissa- and ordinate-values discussed will impart system-specificity, whence combined; (Step 12; **Figure 1**),

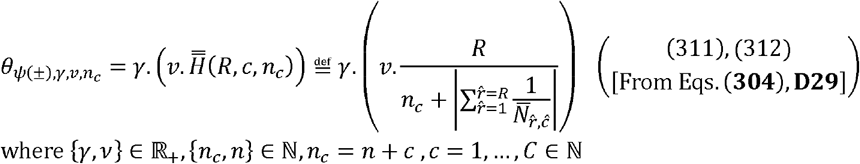

Unperturbed biochemical systems are non-linear and the result of a complex interplay of descriptors many of which include redundancies (molecular, cellular) and remain intangible. Conversely, this implies that a significant percentage of the underlying system (anatomy, physiology) must be altered before the underlying biochemistry is discernible and detectable. In other words, complex biochemical systems encompass degrees of refractoriness and plasticity and are likely to exhibit a switch-like response.

Switching is modeled with activation functions (AFs) and include the rectified linear unit (ReLU)-, *tanh*-, *sigmoid*-, exponential- and softmax-functions amongst several others [74-76]. Although the ReLU-function 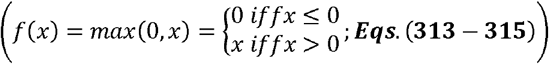 is motivated by neuronal firing and simple, it is an unlikely candidate to recapitulate the inherent complexity of biochemical systems which are non-linear and possess multiple stable states. Whilst modifications such as leaky ReLU 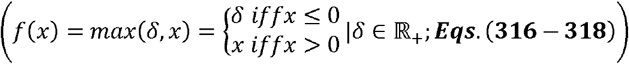 may offset some of these concerns, the unboundedness, limited-choice for the hyperparameter (*δ*) and simplicity, for a complex biochemical system, remain unresolved. Some of these critiques can be addressed by the *sigmoid* - and *tanh*-functions. Whilst the −1, 1-boundedness, positive- and negative-values, steep slope with non-trivial vanishing gradient of the -function are beneficial whence convergence is the desired outcome (classification schema, decision-based causal networks), the same reasons precludes its meaningful deployment as representative of complex biochemical systems [74-76]. Interestingly, the *tanh*-function was purported as a suitable empirical model to represent cooperative binding [75]. In contradistinction, the 0,1-boundedness, strictly positive numerical values, potential scalability and broader slope with a reduced vanishing gradient of the *sigmoid*-function have found favor in representing biochemical systems (accuracy, consistency) [74-76].

Here, we adopt the *sigmoid-*function (*σ* (⋅)) to model cooperative binding for both, catalytic- and non-catalytic systems.

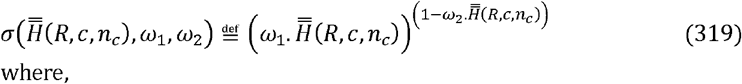

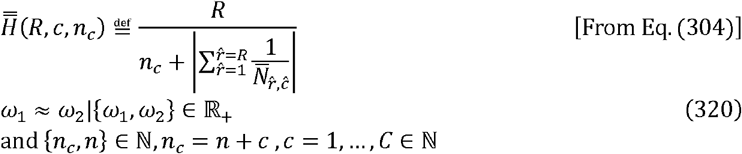

Our discrete-to-continuous model, for the system of ensemble distributions, is a piecewise smooth curve and can be represented bijectively with the sigmoid-function,

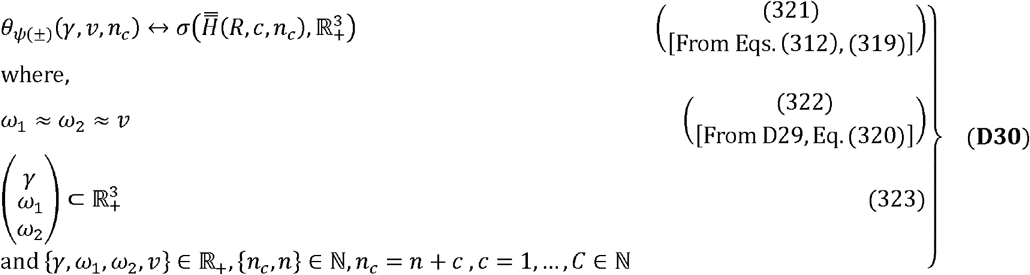

Combining the results, vide supra, we get the explicit equation; (Step 13; **Figure 1**).

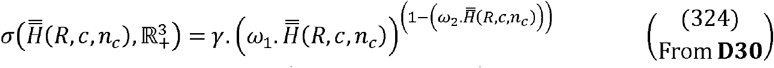

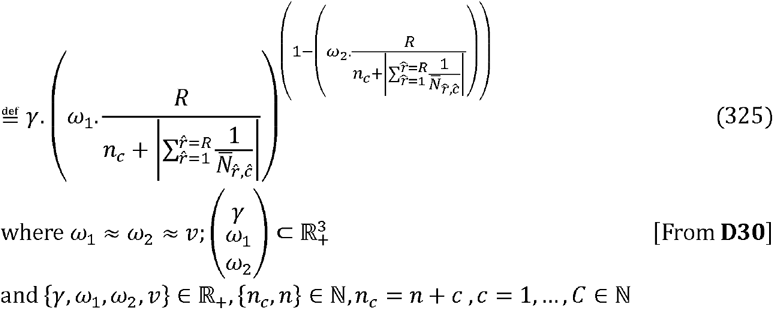

### 4.2 Biochemical relevance of our discrete-to-continuous mathematical model for ensemble distributions of ligand-interacting macromolecular species across milieu-dependent conformational states

We combine our assertions, results and analyses with available biochemical data to highlight the biochemical relevance of our discrete-to-continuous mathematical model for ensemble distributions of ligand-interacting macromolecular species across milieu-dependent conformational states. We prove that our model is phenomenological and is not only able to recapitulate the basic tenets of cooperative deploy the ODC for the Hemoglobin variants (*HbFF, HbAA, HbSS*) and a generic model for the binding but can also assist in gleaning insights into the genesis and progression of the same. We kinetic data generated by the catalytic conversion of L-Aspartate by ATCase, as case studies [75-80].

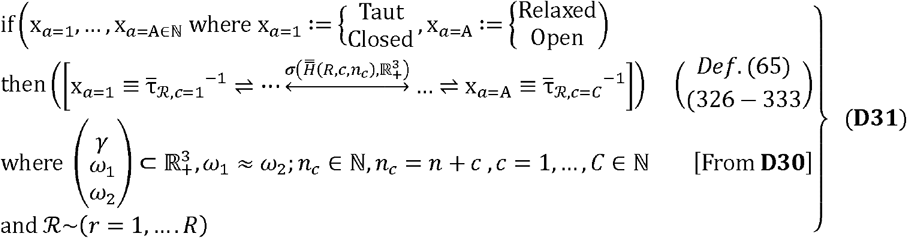

Although cooperative binding, for the case studies presented, is Boolean (closed, open; taut, relaxed), here, we decompose these outcomes into systems of ensemble distributions of distinct milieux-dependent conformational states and model the same in accordance with **D31** and available empirical data wherever applicable (subsections: 4.2.1, 4.2.2)

#### 4.2.1 The oxygen-dissociation curve for Hemoglobin and some of its variants

Hemoglobin (Hb) is characterized by tetrameric (dimer of dimers) functional units for adult Hemoglobin (*HbAA* : = α_2_β_2_| α= 146 aa, β = 110aa) for the semiembryonic fetal hemoglobins (*HbFF* : = {α_2_γ_2,_ ζ_2_β_2_}|α = 146 aa, β = 110aa, ζ = 142 aa, γ = 147 aa) [18, 19]. *HbAA*,, transitions between the alveolar-endothelial interface, peripheral circulation and tissues/organs as well-defined and oxygen-centric conformational states (deoxygenated, partially oxygenated, fully oxygenated),

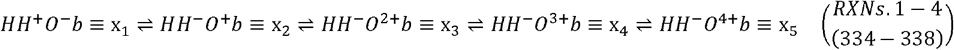

where

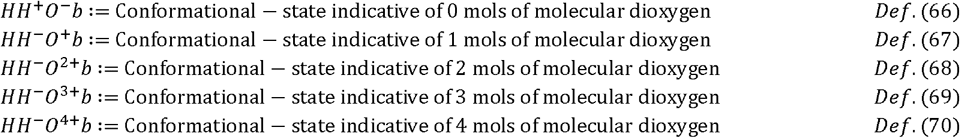

*HbFF*, unlike *HbAA*, interacts poorly with BPG (cavity incongruence, *H*143*S*; structural mismatch, <9.0 Ang), and will therefore, retain most of its bound molecular dioxygen whence at the tissues and is, therefore, better suited to extracting molecular dioxygen from the maternal blood (feto-placental circulation) [18, 19, 75-79]. The *HbSS*-variant, is the Hemoglobin associated with the sickle cell point mutation (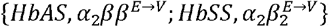) and will attenuate the conformational plasticity needed to achieve a fully oxygenated which, in turn, will lead to a right-shift of the corresponding ODC [18, 19, 75-79].

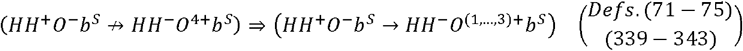

Although there are several parameters to assess the ODC, the *p*50-metric is able to recapitulate the basic tenets of cooperative binding and is defined as the partial pressure of molecular dioxygen when Hemoglobin is 50% saturated and corresponds to the transition between the alveoli (lung; *p*50 = 85 – 90 mmHg; ***Eq***. (**344**)) and peripheral tissues *p*50 = 24 – 28 mmHg; ***Eq***. (**345**)) [18, 19, 75-79]. However, as discussed vide supra, under physiological duress the ODC experiences both, left (lower ambient temperature, higher pH, almost null BPG)- and right (high ambient temperature, low pH, high BPG)-shifts [18, 19, 75-79]. Our presented discrete-to-continuous model is able to model the salient features of the ODC [18, 19, 75-79].

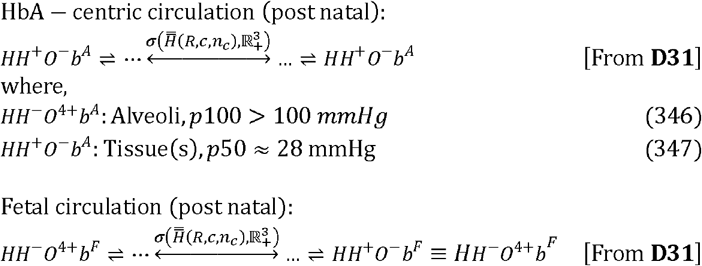

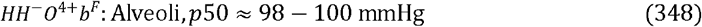

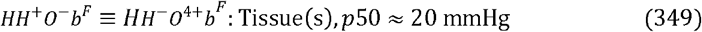

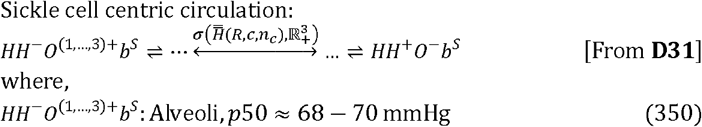

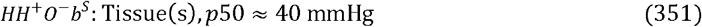

#### 4.2.2 Catalysis by Aspartate transcarbamoylase

The kinetics of binding and thence catalysis, by enzymes, can be altered by small-molecule modifiers such as co-substrates, co-factors (activators, inhibitors) and co-enzymes at one- or more-allosteric sites long-distance conformational changes. Aspartate transcarbomylase (ATCase, EC 2.1.3.2) exhibits cooperative kinetics whence catalyzing the *de novo* synthesis of the pyrimidine nucleus whilst being regulated, allosterically, by cytidine triphosphate (CTP) and adenosine triphosphate (ATP) [41, 42, 62, 80]. We can model these and annotate this as an initial-/terminal-state,

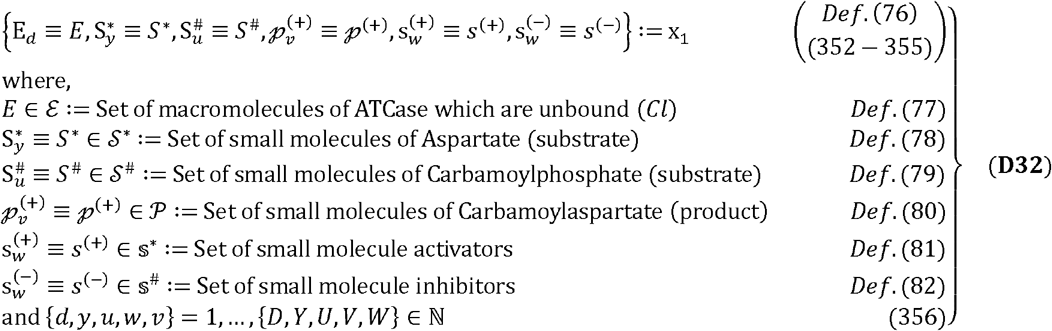

Once catalysis is initiated the following reactions and thence early-states, for ATCase can be appreciated,

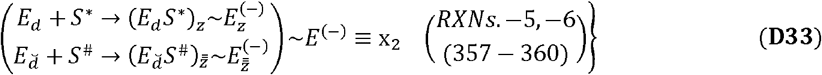

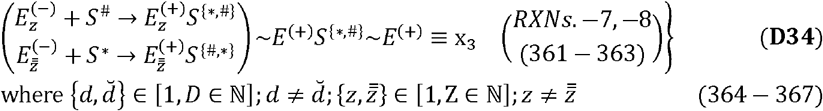

In the presence of ATP, a small molecule activator of ATCase (*s* ^(+)^ : = *ATP*), the enzyme-substrate complex facilitates the formation of Carbamoylaspartate, (*p* ^(+)^),

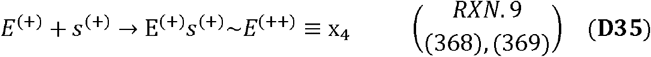

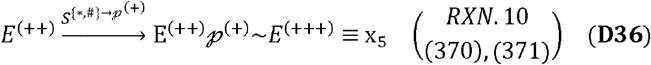

Since Carbamoylaspartate is likely to exhibit sub-optimal binding, to ATCase, it will be released along with ATP whereupon ATCase will revert to the unbound state,

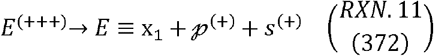

The complete process for positive cooperative binding is then summarized as,

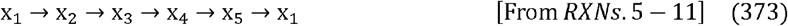

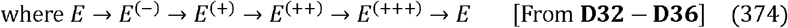

In the presence of CTP, a small molecule inhibitor of ATCase (*s*^(−)^ : = *CTP*), the enzyme-substrate complex facilitates is rendered unstable and dissociates,

On the other hand, if an allosteric inhibitor *s*^(−)^ of ATCase such as Cytidine triphosphate (CTP) is introduced,

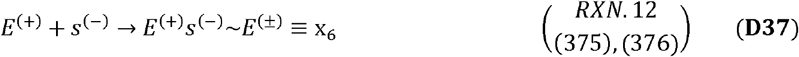

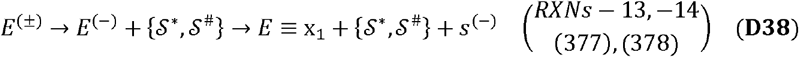

Negative cooperativity, for ATCase, can also be easily demonstrated with our discrete-to-continuous model,

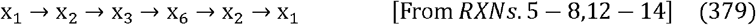

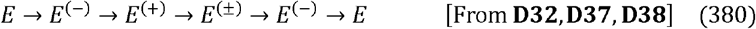

Our discrete-to-continuous model is able to represent both, positive (activator)- and negative (inhibitor)-cooperative binding (**D32**-**D38**) [41, 42, 62, 80].

## 5 Concluding remarks

The kinetics of cooperative binding/unbinding offers insights into ligand-macromolecular interactions, complex formation/disassembly and subtle structural descriptors that are likely to impact biochemical function. Our discrete-to-continuous mathematical model, for ensemble distributions of a ligand-interacting identical macromolecular species across milieux-dependent conformational states, is a piecewise smooth curve (end-end compact set with regular- and smooth-intervals) and is mapped to the sigmoid function. Our model is phenomenological, rather than empirical, and can readily be parameterized with coefficients that are empirically relevant. Our discrete-to-continuous mathematical model is able to recapitulate the fundamental tenets of cooperative binding whilst offering insights into the genesis and progressions of cooperative binding. Our model is scientifically sound, the constructing algorithm is mathematically rigorous, assumptions are unbiased, assertions are biochemically relevant and is able to offer plausible explanations into empirically observed instances of cooperative binding, both enzymatic and non-enzymatic. Future work will consider the effect of heterotypic ligands and/or the effects of finite numbers of allosteric sites, additional- and/or alternate-parameters, explore alternate solutions or functions to approximate empirical data.

## 6 Detailed proofs of a discrete-to-continuous model, for ensemble distributions of a ligand-interacting macromolecular species across milieux-dependent conformational states, of cooperative binding

### Proof (L1)

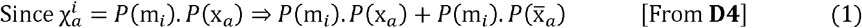

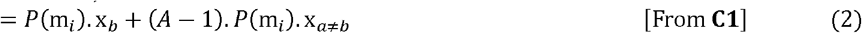

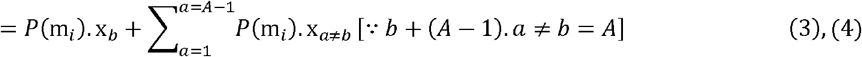

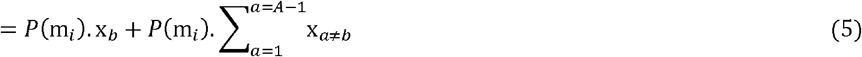

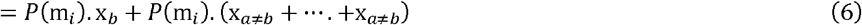

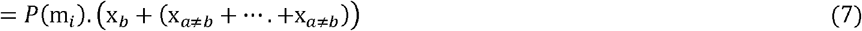

Writing Eq. (7) as a finite series in *a* s.t 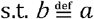

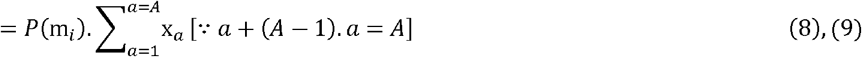

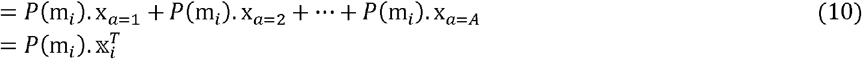

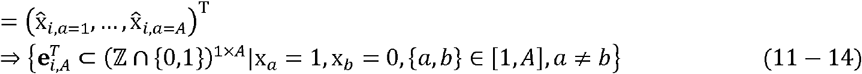

### Proof **C2**

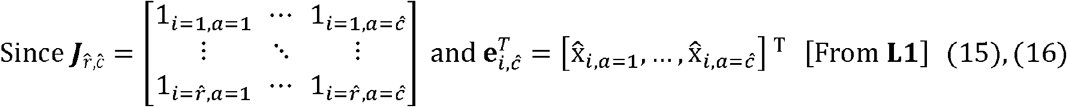

It follows,

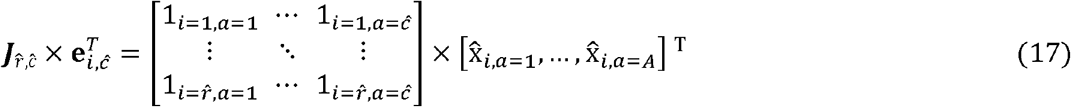

Extending this for ∀*i* and rewriting Eq. (17)

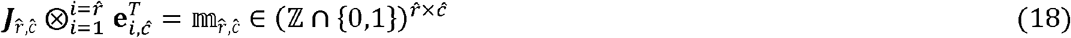

where,

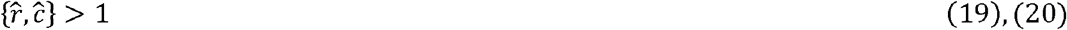

### Proof (**T1**)

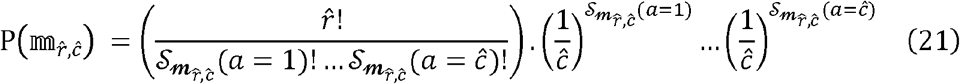

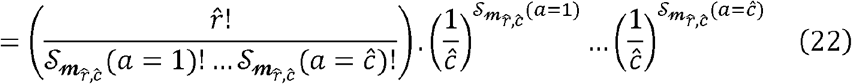

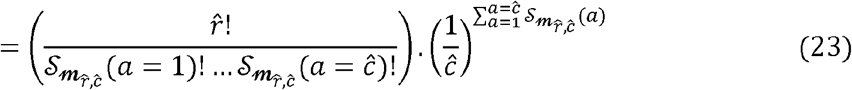

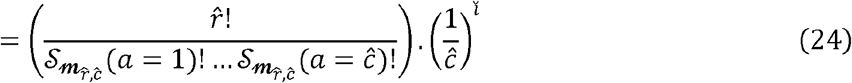

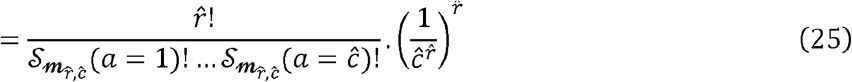

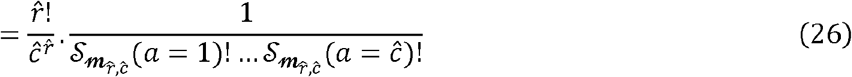

where,

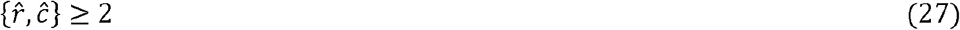

### Proof (**L4**)

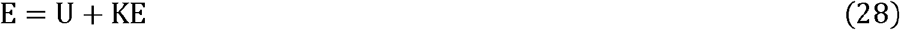

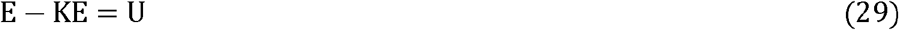

Taking the natural logarithm for both sides

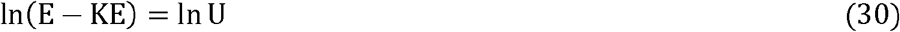

Rewriting LHS in terms of thermodynamic annotation

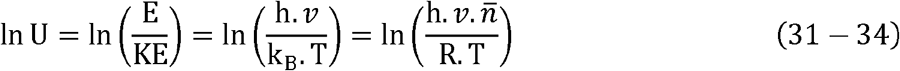

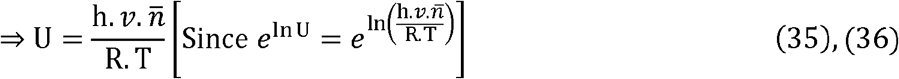

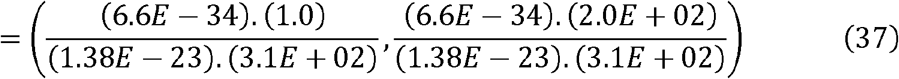

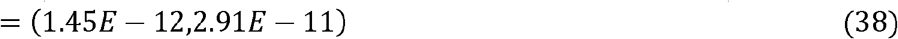

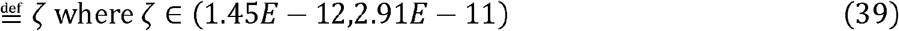

### Proof (C9)

**(C9.1):** Comparing the potential energies for a k, I-pair of ensembles,

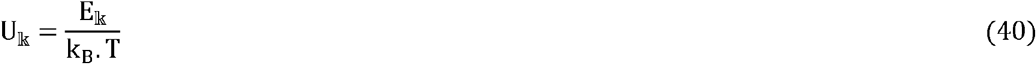

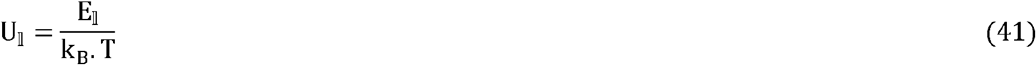

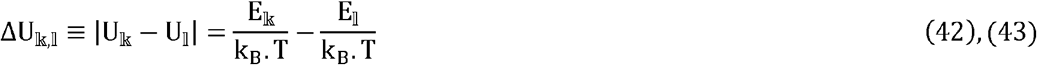

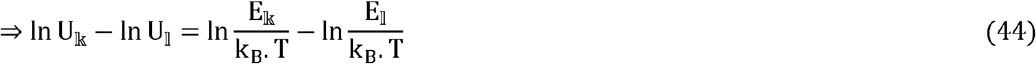

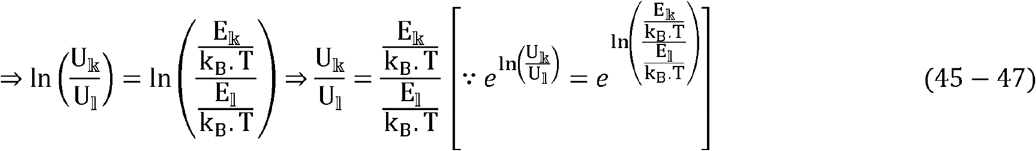

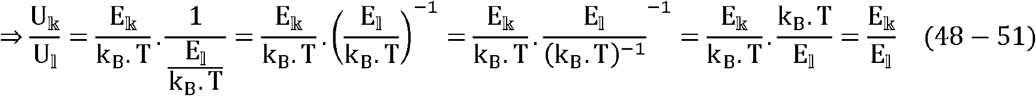

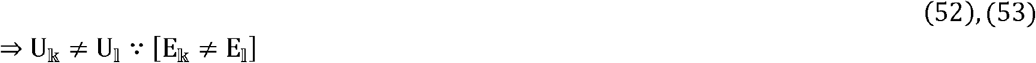

(**C9.2**) Comparing the theoretical probabilities for a distinct k, I-pair of ensembles,

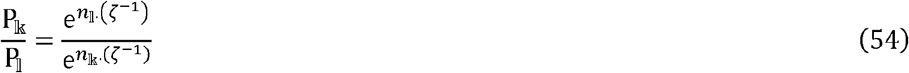

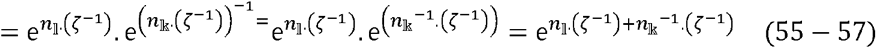

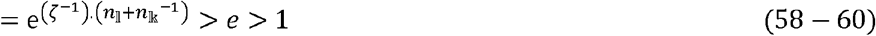

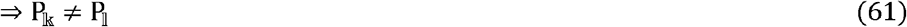

### Proof (**C10**)

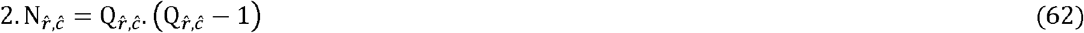

Rearranging and rewriting (70) we get a quadratic equation in 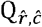,

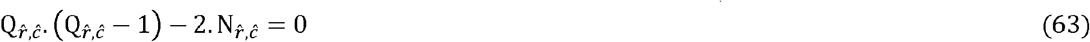

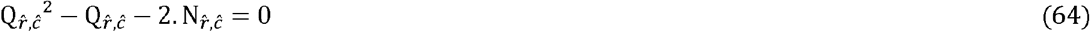

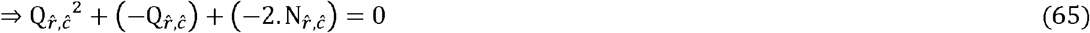

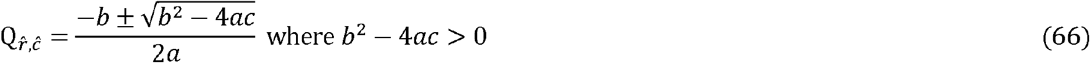

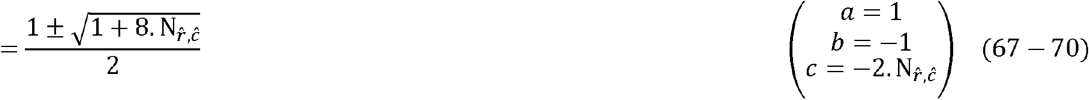

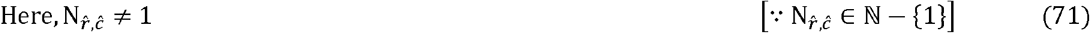

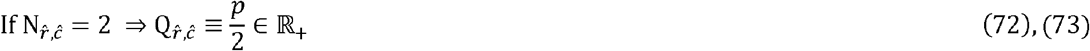

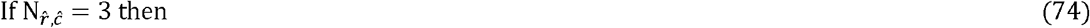

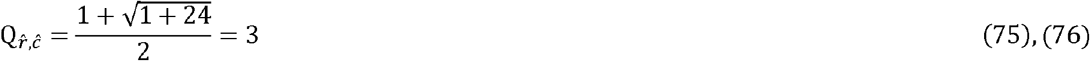

### Proof (**L6**)

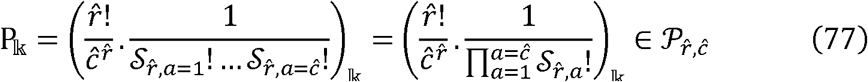

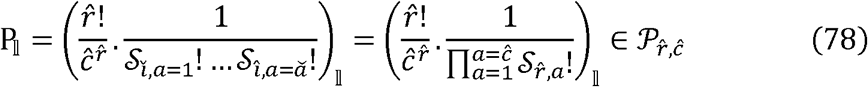

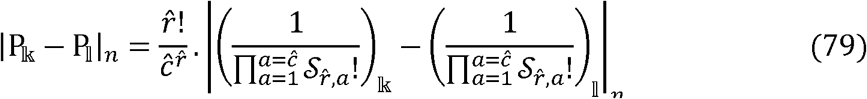

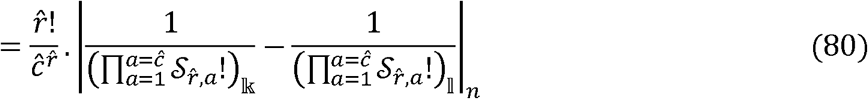

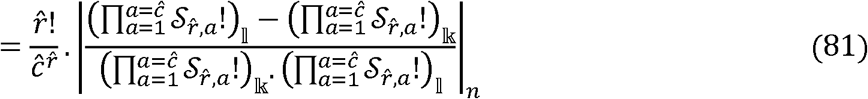

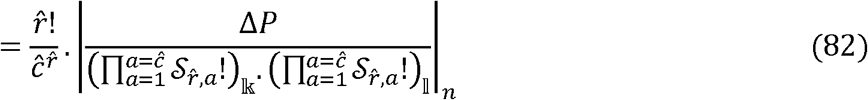

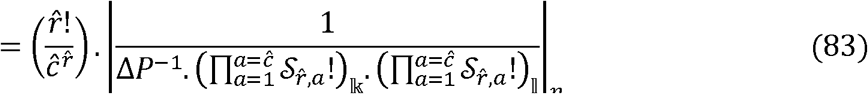

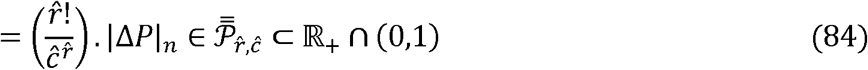

where,

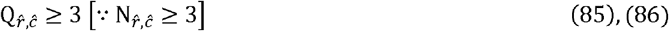

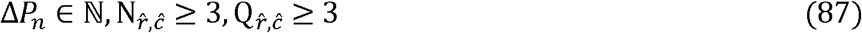

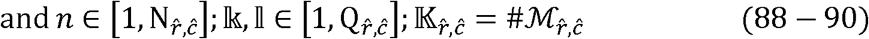

### Proof **L8**

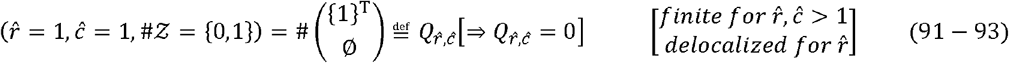

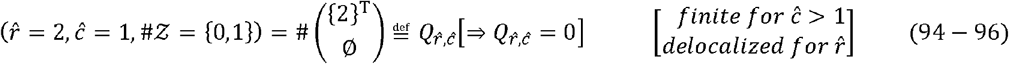

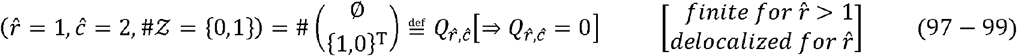

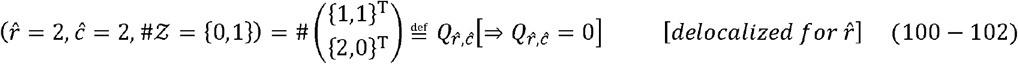

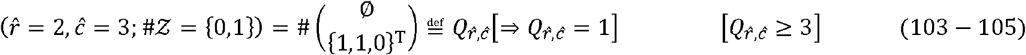

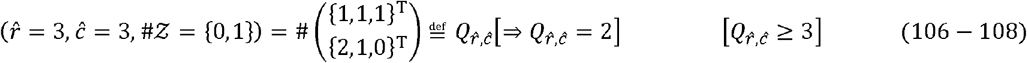

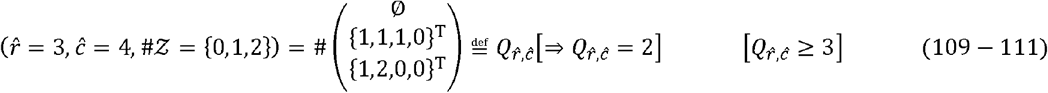

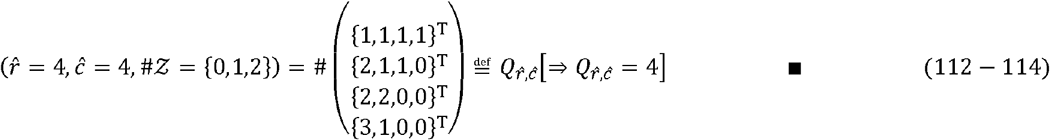

### Proof **T4:**

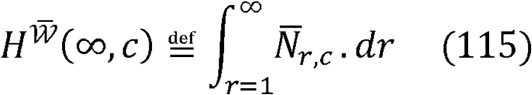

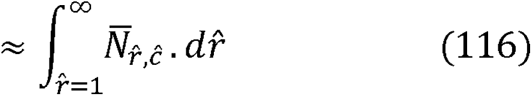

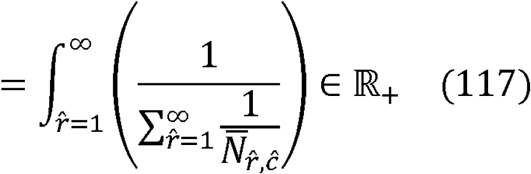

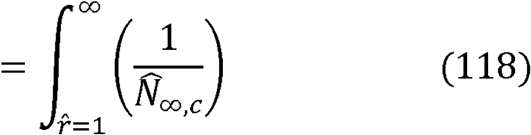

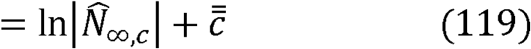

where,

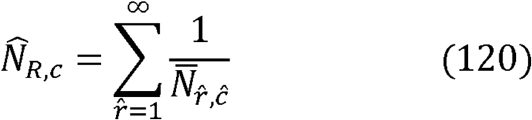

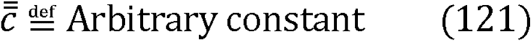

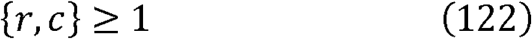

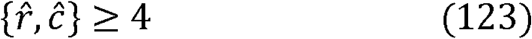

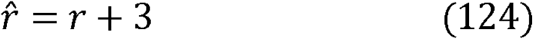

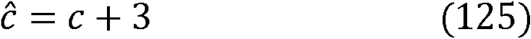

### Proof (L11)

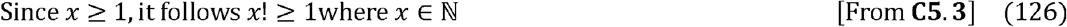

Since *y* >. 1, we can infer the existence of the non − decreasing sequence [From **C5.3**]

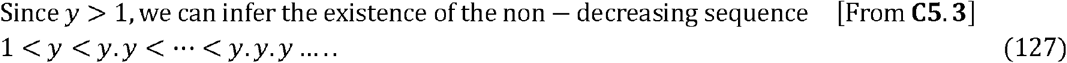

Rewriting this sequence in terms of the exponents of *y*

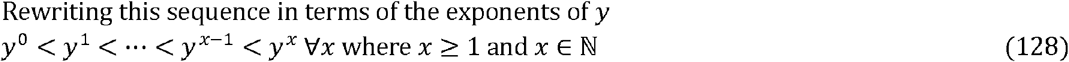

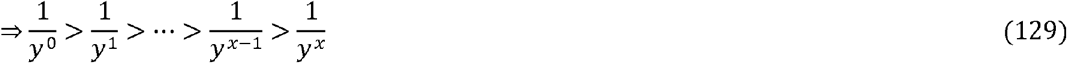

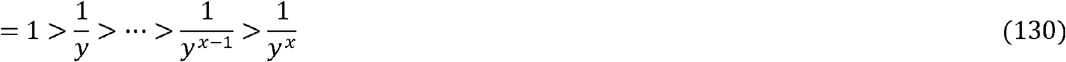

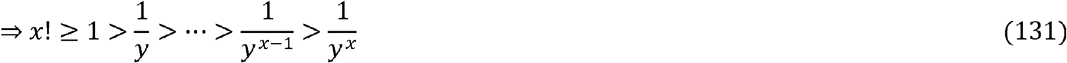

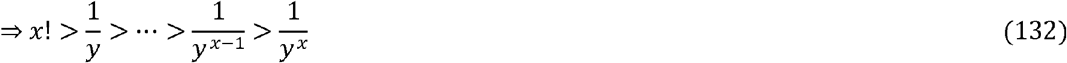

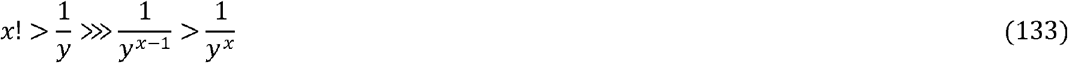

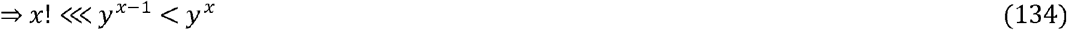

From **C5.3** and **Eq**.(**134**) it follows,

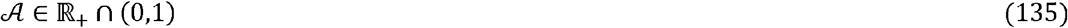

### Proof (T5)

Rewriting 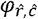 terms of *r,c*

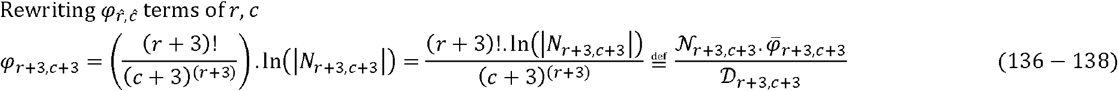

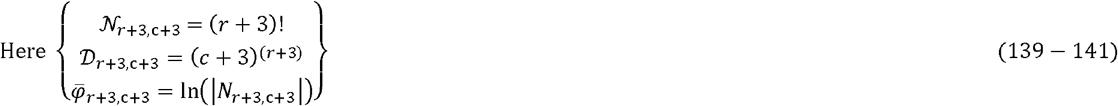

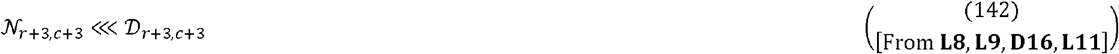

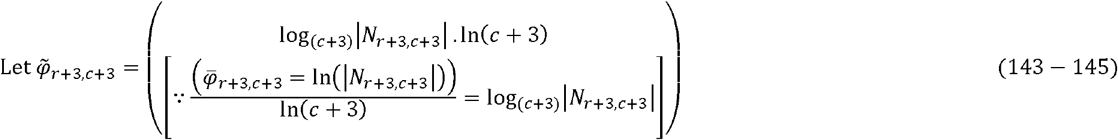

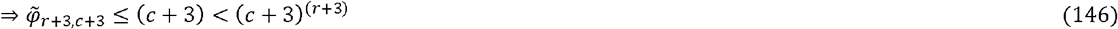

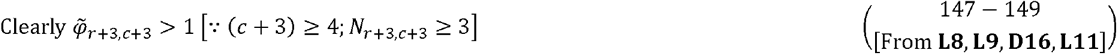

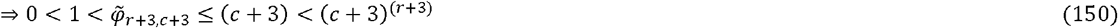

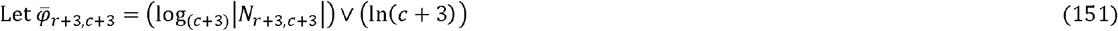

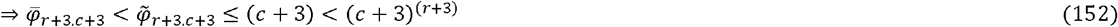

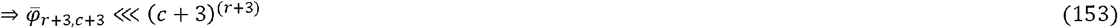

Combining Eqs. (**142**) and (**153**)

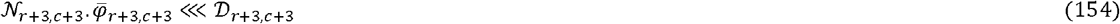

From Eqs. (**150**) and (**154**) it follows,

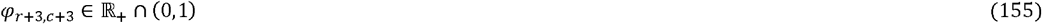

where,

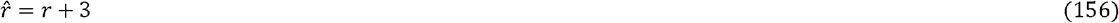

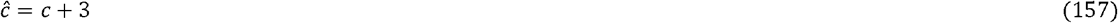

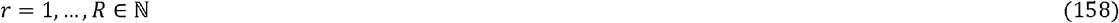

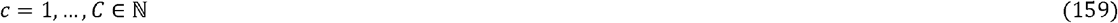

### Proof (C14)

[From **L8, L9, D16, L11, T5**],

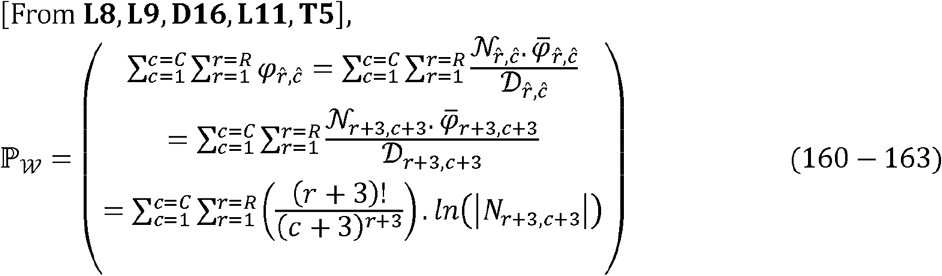

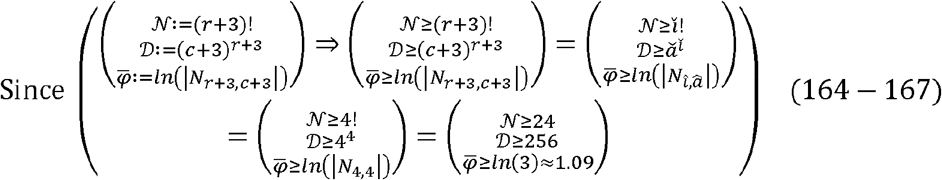

It follows,

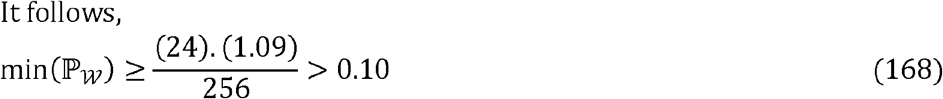

where,

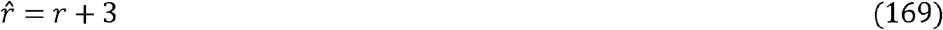

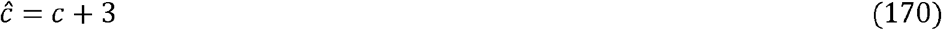

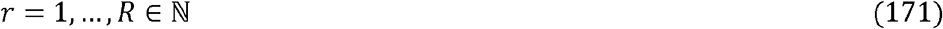

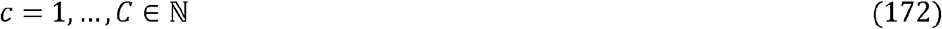

### Proof (**T6)**

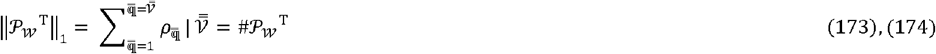

Rewriting Eq. (**173**) as a sequence of partial sums

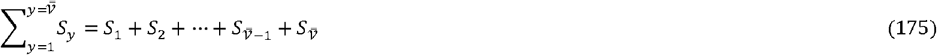

where,

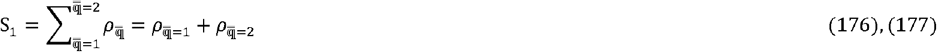

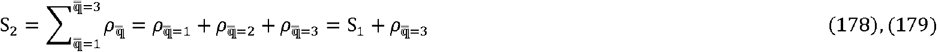

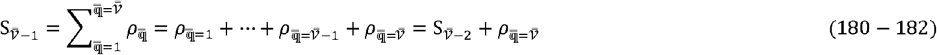

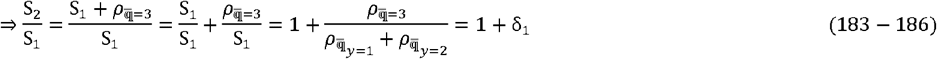

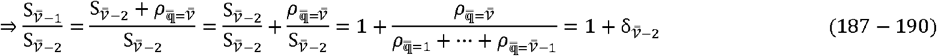

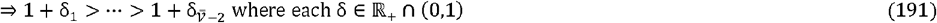

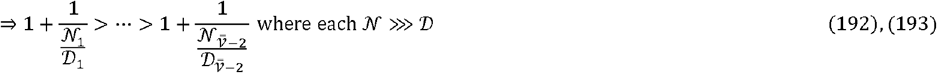

These inequations can be rewritten as,

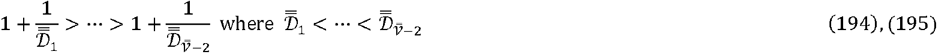

Taking the natural log for each 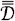

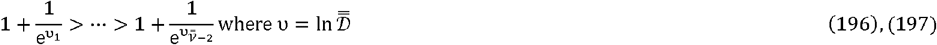

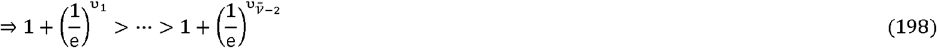

Rewriting these as equations of binomial expressions with integer exponents

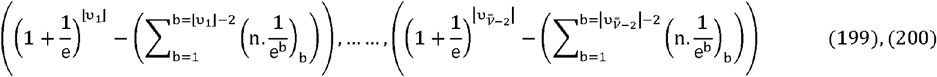

Where n, b ∈ ℕ

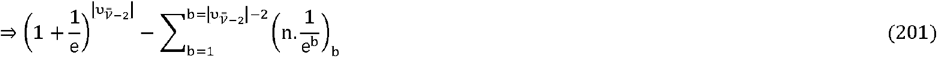

Taking limits

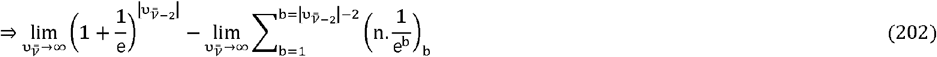

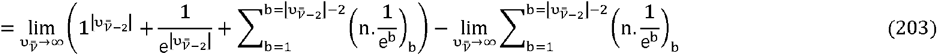

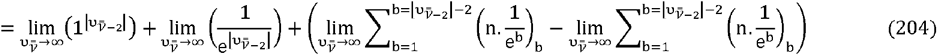

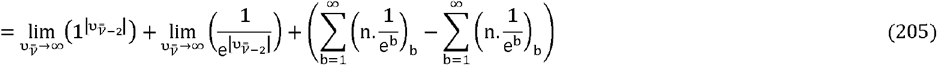

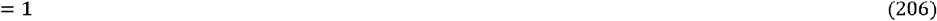

### Proof (T7)

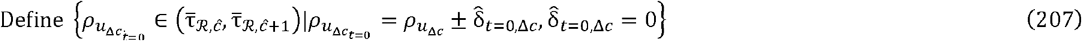

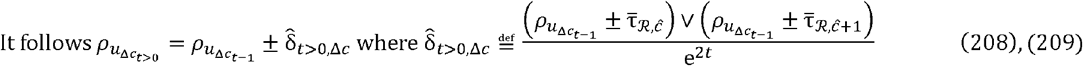

Case 1 (*x* = 1, …, *T* ∈ ℕ):

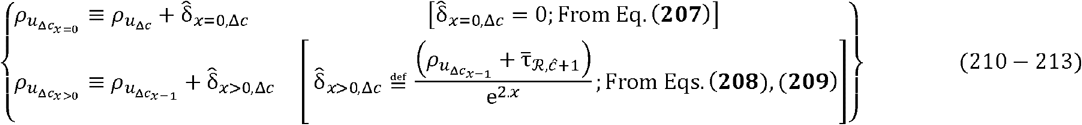

As *x* = 1,2 … *T* it follows,

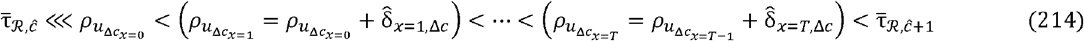

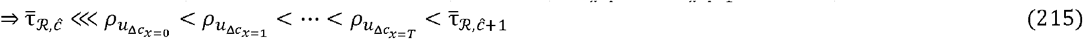

Case 2 (*y* = 1,2 … *T* ∈ ℕ):

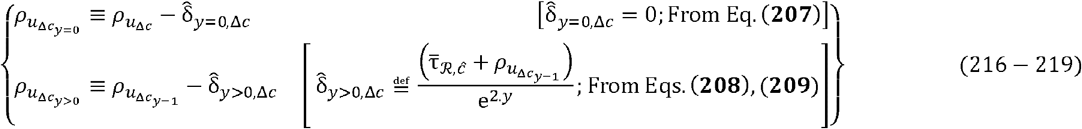

As *y* = 1,2, … *T* it follows,

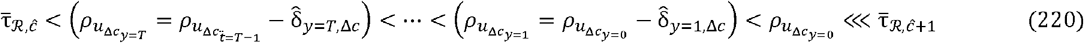

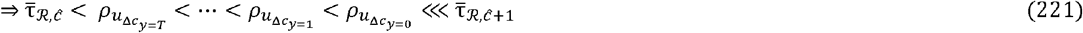

Combining Eqs. (**215**),(**221**)

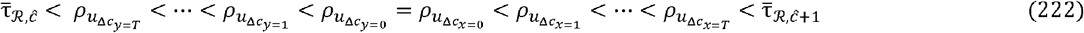

We now reannotate all 2.T + 1 terms associated with any 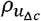, [From Eq. (**222**)]

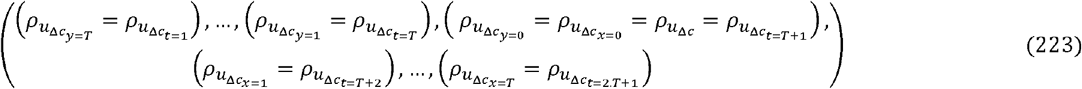

Combining all 2.T + 1 terms associated with any interval − specific 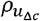 [From Eq. (**223**)]

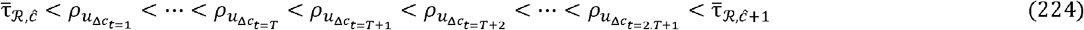

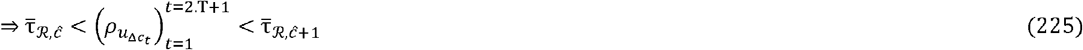

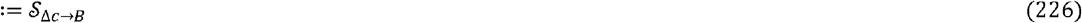

For each term of Eq. (**224**) it follows,

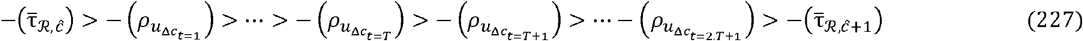

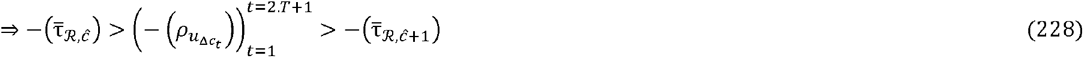

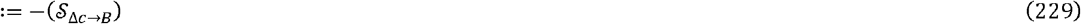

From(**226**), (**229**) it follows

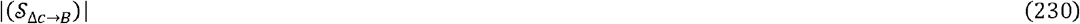

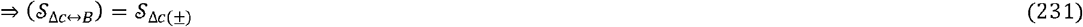

## Acknowledgements

Sk did not receive funding for this work.

## Author Contributions

SK, designed the study; defined, formulated, developed and wrote proofs for the theorems, lemmas and corollaries; developed the assessment metrics and algorithms; conducted numerical studies and collated data; wrote all necessary code and manuscript.

